# Aging increases the distinctiveness of emotional brain states across rumination, worry, and positive thinking

**DOI:** 10.1101/2024.10.29.620853

**Authors:** Masaya Misaki, Aki Tsuchiyagaito, Salvador Guinjoan, Martin Paulus

## Abstract

Emotional well-being improves with age, but how neural activity related to emotional brain states changes with aging remains unclear. This study examined individualized brain activation patterns for rumination, worry, and positive thinking to explore age-related variations in emotional state representation. Thirty-five participants (aged 18-64) recalled autobiographical events tied to these emotional states during fMRI scanning. Brain activity was analyzed using an individualized machine learning classifier. Results showed increased discriminability of rumination and worry with age, with older adults exhibiting heightened activation in cognitive control regions during rumination and reduced activation in the cingulate and temporoparietal junction during worry. No significant age-related changes were found for positive thinking, although increased discriminability between positive and negative states correlated with well-being (FDR < 0.05). These findings suggest that aging enhances cognitive control during rumination and reduces anxiety responses during worry, potentially contributing to improved emotional well-being in older adults.

## Introduction

Emotional processing evolves across the lifespan with emotional well-being generally improving as individuals age ^1,2^. Older adults often report higher levels of well-being compared to younger adults ^3^, which is linked to a decrease in negative thinking with age ^4^. Among negative emotional states, repetitive negative thinking (RNT) is particularly pronounced and significantly impacts mental health. Characterized by persistent, recurring thoughts focused on negative past experiences or future concerns, RNT is associated with an increased risk of anxiety and depression ^5^. Beyond mental health, RNT affects overall cognitive functioning, contributing to executive function deficits in young adults ^6^ and increasing the risk of Alzheimer’s disease in older adults ^7,8^.

Rumination and worry are the main components of RNT, distinguished primarily by their temporal focus: rumination centers on past negative experiences, while worry involves future uncertainties ^9–11^. Despite their distinct temporal orientations, these processes share cognitive features, prompting debates about their conceptual overlap and differences. Psychological assessments have shown that rumination and worry often share meta-cognitive characteristics and appraisal dimensions beyond their temporal focus ^12–14^. Factor analyses suggest that rumination and worry frequently load onto a common factor, although they also retain unique elements ^15,16^. In clinical settings, both processes are correlated but have distinct impacts: rumination is more predictive of depressive comorbidity and recurrence, while worry is closely linked to anxiety disorders ^17,18^.

Given that rumination and worry differ primarily in their temporal focus, age-related variations in how these thought processes manifest may be expected. Indeed, a study examining the effect of age on the valence of ruminative thoughts found that negative ruminative thoughts decreased in older adults, while there was no significant age-related difference in positive repetitive thoughts ^19^. Since RNT often centers on self-relevant events ^20^ and emotional responses change with age ^1,2^, understanding individual differences in the neural representation of these emotional states is crucial. These differences could provide insights into the distinctiveness of these cognitive processes and their impact on emotional well-being across the lifespan.

In this study, we investigated the brain processes underlying rumination, worry, and positive thinking in non-clinical individuals, examining how these emotional states are represented in individualized brain activation patterns and how these patterns vary with age. Brain activation was analyzed during thoughts about past negative events (rumination), future concerns (worry), and positive experiences. Emotional states are increasingly understood to be reflected in whole-brain activity patterns rather than isolated brain regions ^21–24^. Aligning with this holistic view of brain function, we hypothesized that these emotional states would be distinguishable by their overall brain activity patterns rather than by specific regional activations. To this end, we employed machine learning models to identify distinctions between these states that may not be detectable through traditional mass voxel-wise analysis.

Additionally, we built personalized models to capture individual-specific neural representations of these cognitive processes. Given that RNTs are centered on self-relevant events ^20^, neural representations of these emotional processes are expected to become more distinct when individuals reflect on personal experiences ^25^. Moreover, common discrimination models often carry biases toward stereotypical profiles ^26^, potentially obscuring subtle differences between individuals. By using personalized models, we aimed to minimize these biases and provide a more accurate evaluation of individual-specific brain activity patterns associated with these emotional states.

We examined how the distinctiveness of rumination, worry, and positive thinking relates to demographic factors, particularly age, as well as sex, and measures of rumination, anxiety, and positive affect. This study aims to uncover how aging shapes the neural representation of these emotional and cognitive states, offering insights into the evolution of emotional brain states across the lifespan. Our findings contribute to a more nuanced understanding of how age influences emotional well-being and cognitive processing in the brain.

## Results

### Participants and Data Quality

Thirty-seven healthy individuals (24 females, aged 18–64) participated in the study. Due to excessive head motion leading to the exclusion of more than two of six runs (see the Methods for the exclusion criterion), two participants were removed from the analysis. Thus, the final dataset included 35 participants. Table 1 presents the demographic and assessment statistics of the included participants.

**Table 1.** Participant demographics and assessment scores. P-values for the age association are uncorrected.

|  |  | Age association |
| --- | --- | --- |
| Sex (n = female/male) | 24/11 | $t = 1.93, p = 0.068$ |
| Age (mean [SD]) | 40.7 [14.0] |  |
| RRS | 31.6 [7.9] | $r = -0.229, p = 0.186$ |
| BSRI | 172.4 [141.3] | $r = -0.348, p = 0.040$ |
| STAI-T | 31.3 [11.0] | $r = -0.328, p = 0.055$ |
| STAI-S | 29.8 [7.9] | $r = -0.362, p = 0.032$ |
| SCI-QOL Positive | 57.1 [7.9] | $r = 0.054, p = 0.758$ |

RRS, Ruminative Response Style; BSRI, Brief State Rumination Inventory; STAI-T/S, State-Trait Anxiety Inventory Trait/State scores; SCI-QOL Positive, Spinal Cord Injury – Quality of life index positive affect and well-being score. $r$ , Spearman correlation.

### Thought State Classification

The participants performed a thought induction task, which included rumination, worry, and positive thinking blocks interlaced with an attentional (flanker) task and resting blocks (Figure 1). The thought state was classified at each time point in the blocks using a machine learning model with inputs of preprocessed fMRI signals in whole-brain gray matter voxels. The model was built for individual participants. This analysis was performed using the Auto-sklearn package ^27^, which constructs ensembles of machine learning models from the scikit-learn library. Model performance was evaluated using the area under the receiver operating characteristic curve (AUC) in a leave-one-run-out cross-validation.

**Figure 1.**
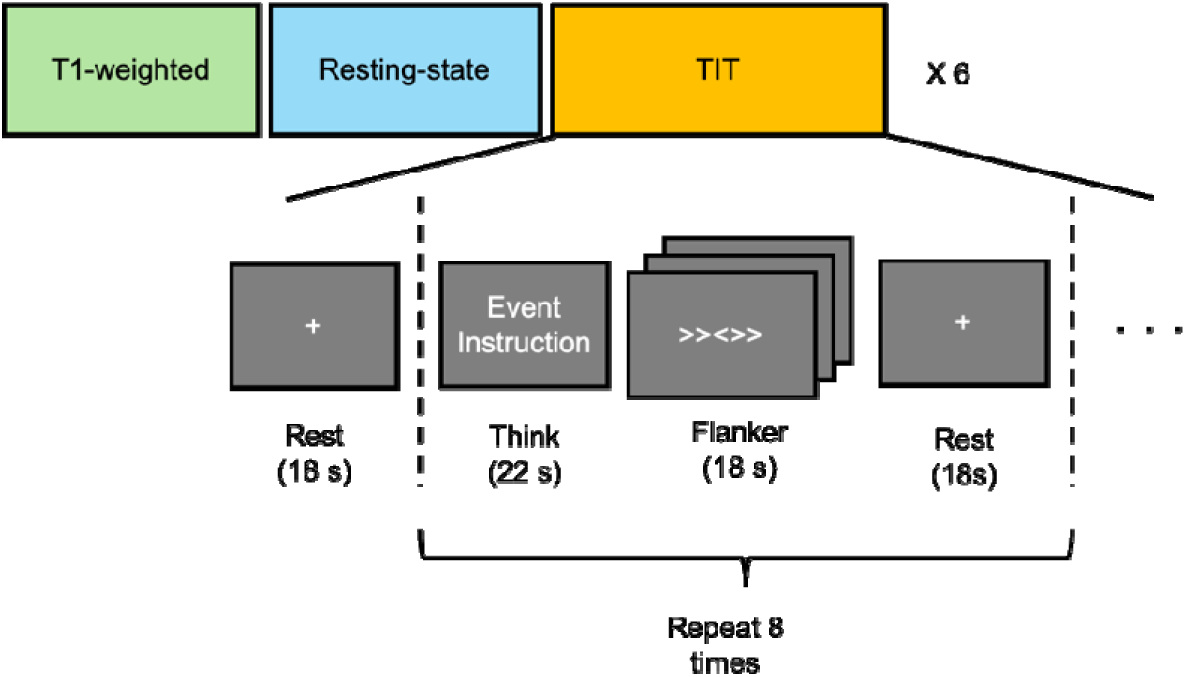
The scan session began with a T1-weighted anatomical scan, followed by a resting- state run (6 min 50 s) and six runs of the TIT. Each TIT run began with an 18 s Rest block, followed by blocks of 22 s Think, 18 s Flanker task, and 18 s Rest. These blocks were repeated eight times per run, with each run lasting 8 min 2 s.

The resting state was clearly distinguishable from the three thinking states (rumination, worry, and positive thinking) in the whole-brain machine learning analysis. All participants demonstrated significant AUC values (*p* < 0.05 with Bonferroni correction) for distinguishing rest from rumination (mean ± SD AUC = 0.879 ± 0.041), worry (0.882 ± 0.048), and positive thinking (0.893 ± 0.059). However, discrimination between the three thinking states varied across individuals. Figure 2 displays the distribution of AUC values for the distinctions between rumination, worry, and positive thinking. The lowest discrimination score was observed between rumination and worry (AUC = 0.648 ± 0.115), with nine participants showing non-significant discrimination. The distinction between rumination and positive thinking (AUC = 0.739 ± 0.116) and between worry and positive thinking (AUC = 0.736 ± 0.117) also showed variability, with four and three participants, respectively, showing non-significant discrimination. These AUC values were not correlated with self-reported measures of task difficulty, successful thinking, sleepiness, emotional exhaustion, or tiredness (see Supplementary Table S1), indicating that the variability in classification was not related to task compliance. Furthermore, AUC values were not associated with distress levels related to rumination or worry events (Supplementary Table S1), suggesting that the discriminability difference was not associated with individual variability in the severity of negative events.

**Figure 2.**
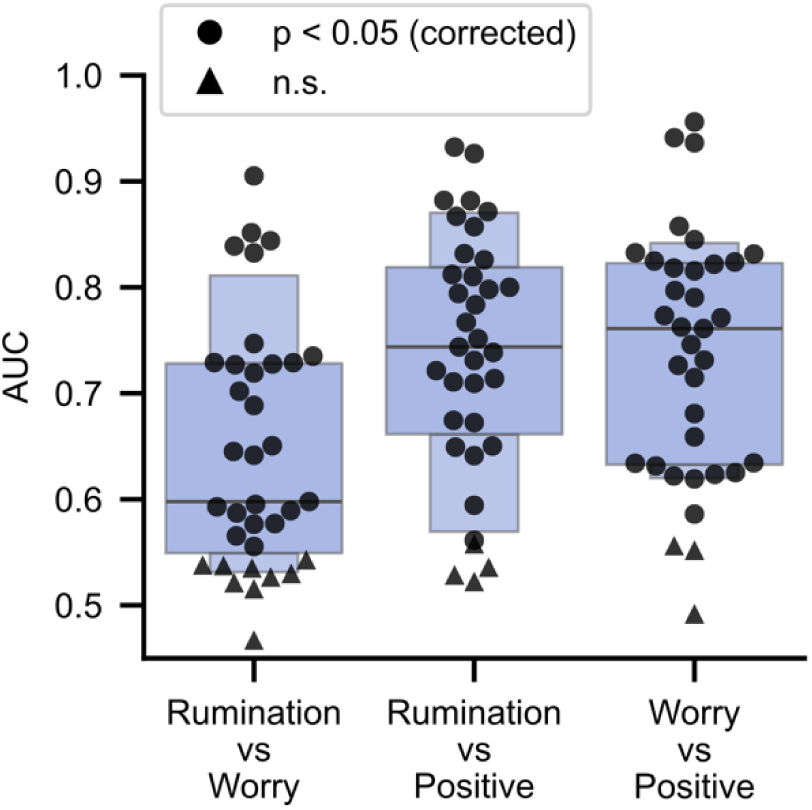
Distribution of AUC values across participants. AUCs from the whole-brain machine learning classification results are shown for each contrast among the three thinking states: rumination, worry, and positive events. The symbols represent the significance of AUC for individual participants. n.s., non significant.

### Individual Differences in Thought State Discrimination

To investigate factors contributing to individual differences in thought state discrimination, we analyzed correlations between AUC values and variables such as age, sex, and assessment scores (Table 2, Figure 3). Age showed a significant correlation with the discrimination of all three states—rumination, worry, and positive thinking—with older participants demonstrating greater discrimination across these emotional states (Figure 3). Additionally, discrimination between negative (rumination and worry) and positive states was significantly correlated with trait rumination (RRS ^28^), trait anxiety (STAI-T ^29^), and positive affect (SCI-QOL Positive ^30^).

**Figure 3.**
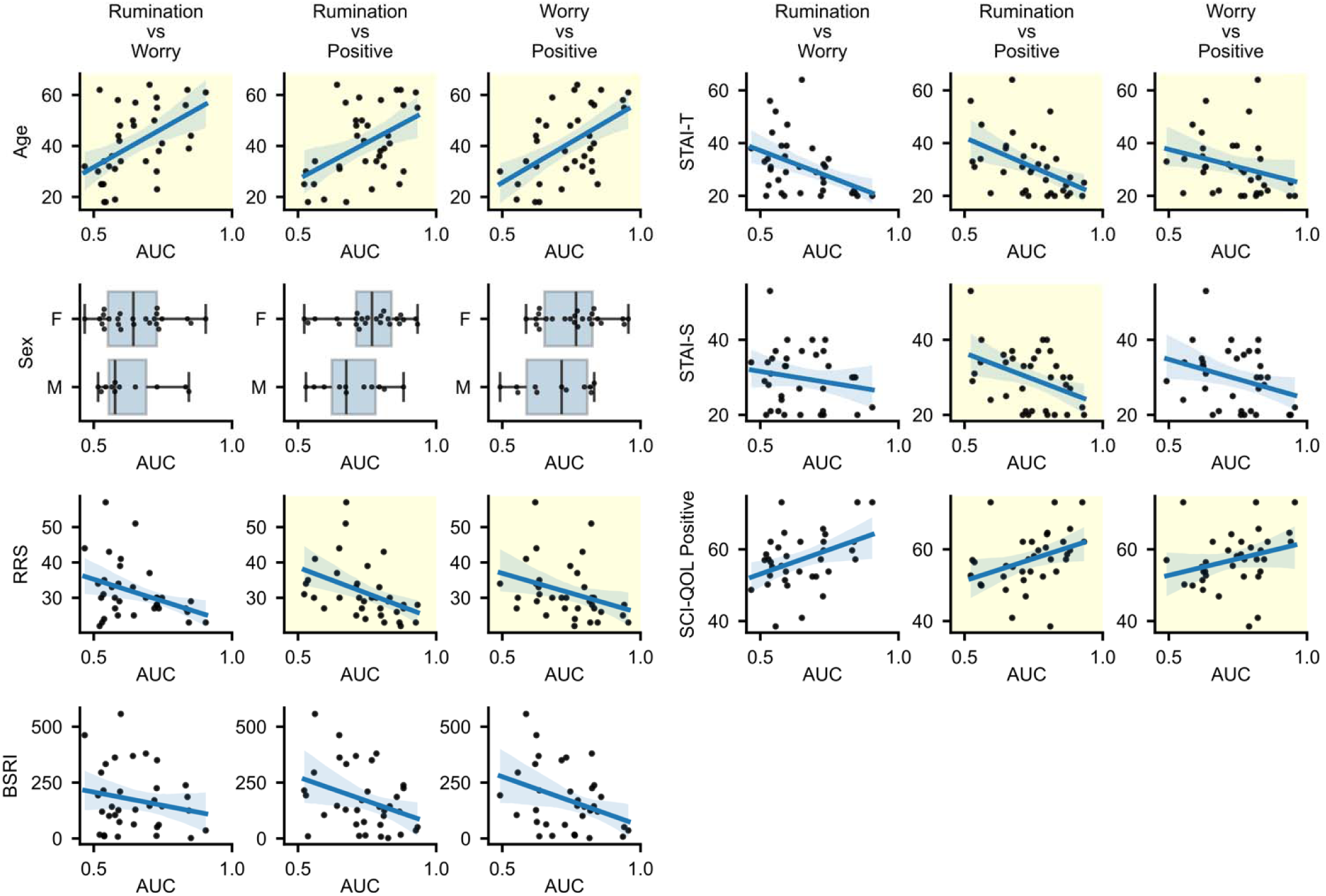
Associations between thought state discrimination (AUC) and participants’ demographic variables and assessment scores. The line represents the fitted linear association, with the shaded area indicating the 95% confidence interval. A yellow background highlights combinations with significant associations (FDR < 0.05, see Table 2). RRS, Ruminative Response Style; BSRI, Brief State Rumination Inventory; STAI-T/S, State-Trait Anxiety Inventory Trait/State scores; SCI-QOL Positive affect, Spinal Cord Injury – Quality of life index positive affect and well-being score.

**Table 2.** Correlations between thought state discrimination (AUC) and participants’ demographic variables and assessment scores. Statistical (stat) values are represented by t-values for Sex and Spearman correlations for all other variables.

|  | Rumination-Worry |  |  | Rumination-Positive |  |  | Worry-Positive |  |  |
| --- | --- | --- | --- | --- | --- | --- | --- | --- | --- |
| | stat | $p$ | FDR | stat | $p$ | FDR | stat | $p$ | FDR |
| Sex | 0.527 | 0.605 | 0.628 | 1.461 | 0.160 | 0.160 | 1.669 | 0.113 | 0.113 |
| Age | 0.494 | 0.003 | 0.018* | 0.410 | 0.014 | 0.025* | 0.485 | 0.003 | 0.017* |
| RRS | -0.303 | 0.076 | 0.134 | -0.589 | 0.000 | 0.001* | -0.466 | 0.005 | 0.017* |
| BSRI | -0.123 | 0.481 | 0.628 | -0.328 | 0.055 | 0.064 | -0.283 | 0.099 | 0.113 |
| STAI-T | -0.380 | 0.025 | 0.086 | -0.517 | 0.001 | 0.004* | -0.406 | 0.016 | 0.036* |
| STAI-S | -0.085 | 0.628 | 0.628 | -0.358 | 0.034 | 0.048* | -0.280 | 0.103 | 0.113 |
| SCI-QOL Positive | 0.352 | 0.038 | 0.088 | 0.514 | 0.002 | 0.004* | 0.383 | 0.023 | 0.041* |

\*, FDR < 0.05; RRS, Ruminative Response Style; BSRI, Brief State Rumination Inventory; STAI-T/S, State-Trait Anxiety Inventory Trait/State scores; SCI-QOL Positive affect, Spinal Cord Injury – Quality of life index positive affect and well-being score.

State anxiety (SATI-S) was specifically correlated with rumination-worry discrimination.

Given that age was the most significant factor overall, we further examined its correlations with other variables (Table 1). Although state rumination (BSRI ^31^) and state anxiety (STAT-S) showed negative correlations with age (uncorrected *p* < 0.05), these relationships were not significant after correction. Additional correlations between AUCs and other exploratory measures are presented in Supplementary Table S2. Results indicated that higher discriminability of the positive state from negative emotional states (rumination and worry) was associated with lower difficulties in emotion regulation (DERS).

### Age-Related Differences in the Time Course of Emotion Emergence

We further examined differences in the time course of decoded states between younger and older participants. Figure 4 presents the mean time course of classifier outputs in young and old participants (divided by median age = 39). Linear mixed-effects (LME) analyses showed a significant main effect of age group for all three states, with older participants exhibiting a higher decoding probability for the corresponding state. However, no significant interaction was found between age group and time for any state, suggesting that age-related differences in thought state discrimination were not linked to variations in the temporal pattern of thought development.

**Figure 4.**
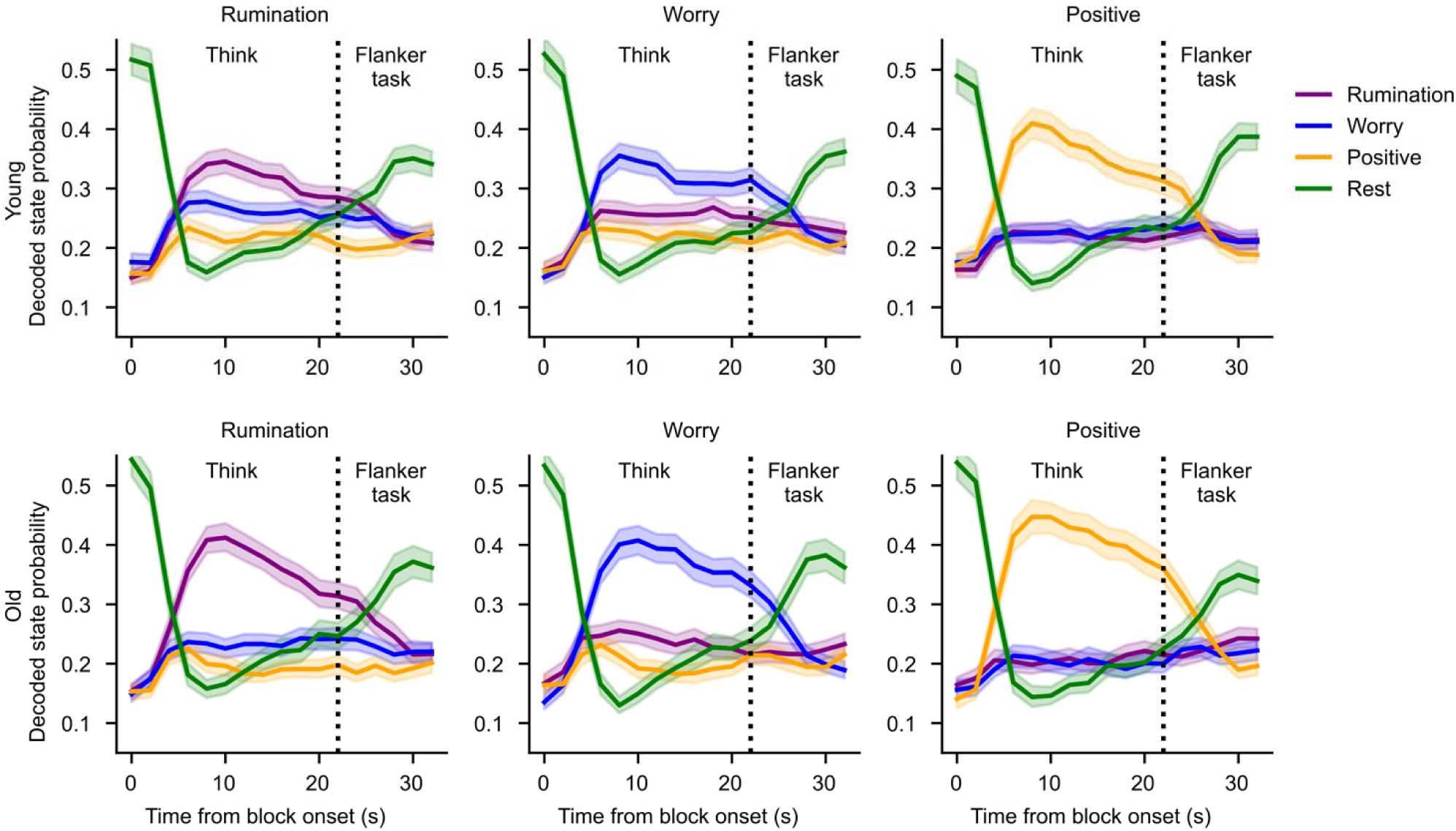
Time course of state probability from the machine learning classifier outputs in each condition of the Think block. Shaded bands represent the 95% confidence interval.

### Brain Activation Patterns for Each Emotional State

We investigated brain activation patterns during each thought state by mapping beta coefficients for the decoded time series of mental state in a general linear model (GLM) analysis (Figure 5). Peak coordinates of significant clusters are detailed in Supplementary Tables S3-S8. All three thought states were associated with activity in the lateral frontal regions, medial premotor and supplementary motor areas (SMA), superior temporal regions, visual cortex, hippocampus, putamen, and cerebellum. Reduced activity was observed in the anterior portion of the posterior cingulate region and the supramarginal gyrus. Both worry and positive thinking states showed positive associations with activation in the precuneus and the posterior part of the posterior cingulate, regions typically linked to self-related thoughts. In rumination, there was more activation in the inferior parietal lobule and less suppression in the supramarginal gyrus, regions also associated with self-referential thoughts, alongside lower activation in the ventromedial prefrontal cortex and visual cortex compared to other states. Worry states showed greater engagement of dorsal frontal regions compared to both rumination and positive thinking. Positive thinking involved higher activation in the retrosplenial cortex, hippocampus, and amygdala.

**Figure 5.**
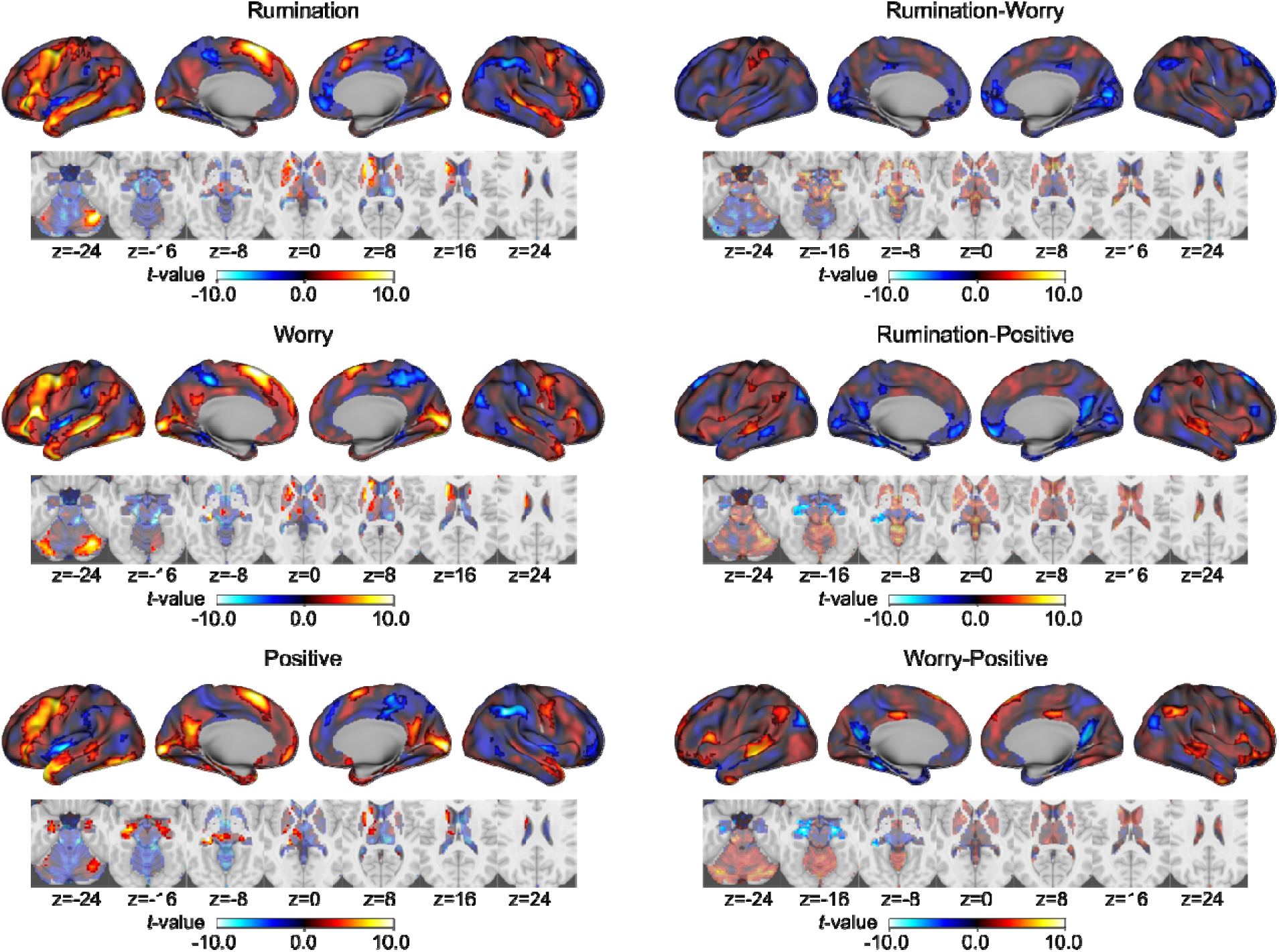
Brain activation patterns associated with the thought states and their contrasts. The maps are displayed on the inflated cortical surface, with subcortical and cerebellar regions are shown on axial slice maps. Clusters with voxel-wise p < 0.001, corrected for cluster extent at p < 0.05, are highlighted with opaque colors. Unthresholded maps are also shown with lower opacity in the plots. The images are displayed in neurological orientation, with the left side corresponding to the left side of the brain.

### Age-Related Effects in Emotional Brain Activation

Figure 6 shows age effects in each thought state and their contrasts. The peak coordinates of significant clusters are provided in Supplementary Tables S9-S13. For the rumination state, age was positively associated with increased activation in broad brain regions, with significant clusters in the right middle frontal gyrus, right superior frontal gyrus, fusiform gyrus, and cerebellum. In contrast, in the worry state, most brain regions showed a negative association with age, with significant clusters observed in the right temporoparietal junction (TPJ) region and anterior and middle cingulate cortex. The positive thinking state did not show a significant association with age.

A contrast of age effects between the rumination and worry states revealed that several key regions associated with self-referential thinking and cognitive control showed greater age-related increases in rumination relative to worry. These include the right middle and inferior frontal gyrus, right supramarginal gyrus, middle temporal and occipital gyri, and the precuneus. The cerebellum and cingulate cortex also exhibited heightened engagement in rumination with advancing age, compared to worry.

When the block-wise response model was used to evaluate brain activations during the think blocks in the GLM, instead of the decoded time series, similar activation patterns were observed for the thought state activations and their contrasts (Supplementary Figure S1).

However, no significant age-related differentiation was found using the block-wise response model.

## Discussion

The brain states of rumination and worry, two core components of RNT, were distinguishable from each other based on whole-brain activation patterns, although the discriminability varied depending on the participant’s age. Older participants exhibited more distinct activation patterns, characterized by increased activation in cognitive control regions during rumination and decreased activation in the TPJ and anterior and middle cingulate cortex during worry. No significant age-related changes were observed in brain activation for the positive thinking state. These results highlight the importance of considering age when investigating the neural underpinnings of emotional states and the differentiation between rumination and worry. While the commonality of these RNT components has been demonstrated in psychological assessments and factor analyses, incorporating a lifespan perspective on these emotional states may yield new insights into the factors driving their differentiation.

The absence of significant age effects on positive thinking-related brain activity observed in this study aligns with previous findings on lifelong changes in emotional states. Older adults often report higher levels of well-being compared to younger adults, which is linked to a reduction in negative thinking as they age ^3,4^. However, this reduction does not extend to positive repetitive thoughts, which remain stable across age ^19^. In the current study, while no significant correlation was found between age and positive affect or well-being scores (SCI-QOL Positive), the separability of brain activity between positive and negative thinking was significantly associated with well-being scores (Table 2). These findings suggest that age-related changes in brain activation patterns during negative thinking may play a key role in enhancing well-being in older adults, supporting the view that modifications in negative emotional processing are crucial for improved emotional well-being with age.

Moreover, the current results showed that rumination and worry were associated with distinct alterations in brain activity with age among negative thought patterns. A significant increase in brain activity was observed during rumination, particularly in regions related to cognitive control, including the lateral frontal and superior parietal areas (Figure 6). Additionally, a greater distinction between rumination and positive thinking was linked to reduced levels of rumination and anxiety (Table 2). These findings suggest that, with age, individuals may respond more adaptively to negative memories by enhancing cognitive control, thereby reducing the rumination and anxiety associated with these memories. This aligns with the suggestion that older adults achieve well-being by optimizing emotion regulation process ^32^ and the view that aging is a process of adaptive change, rather than a decline in cognitive function ^33^.

**Figure 6.**
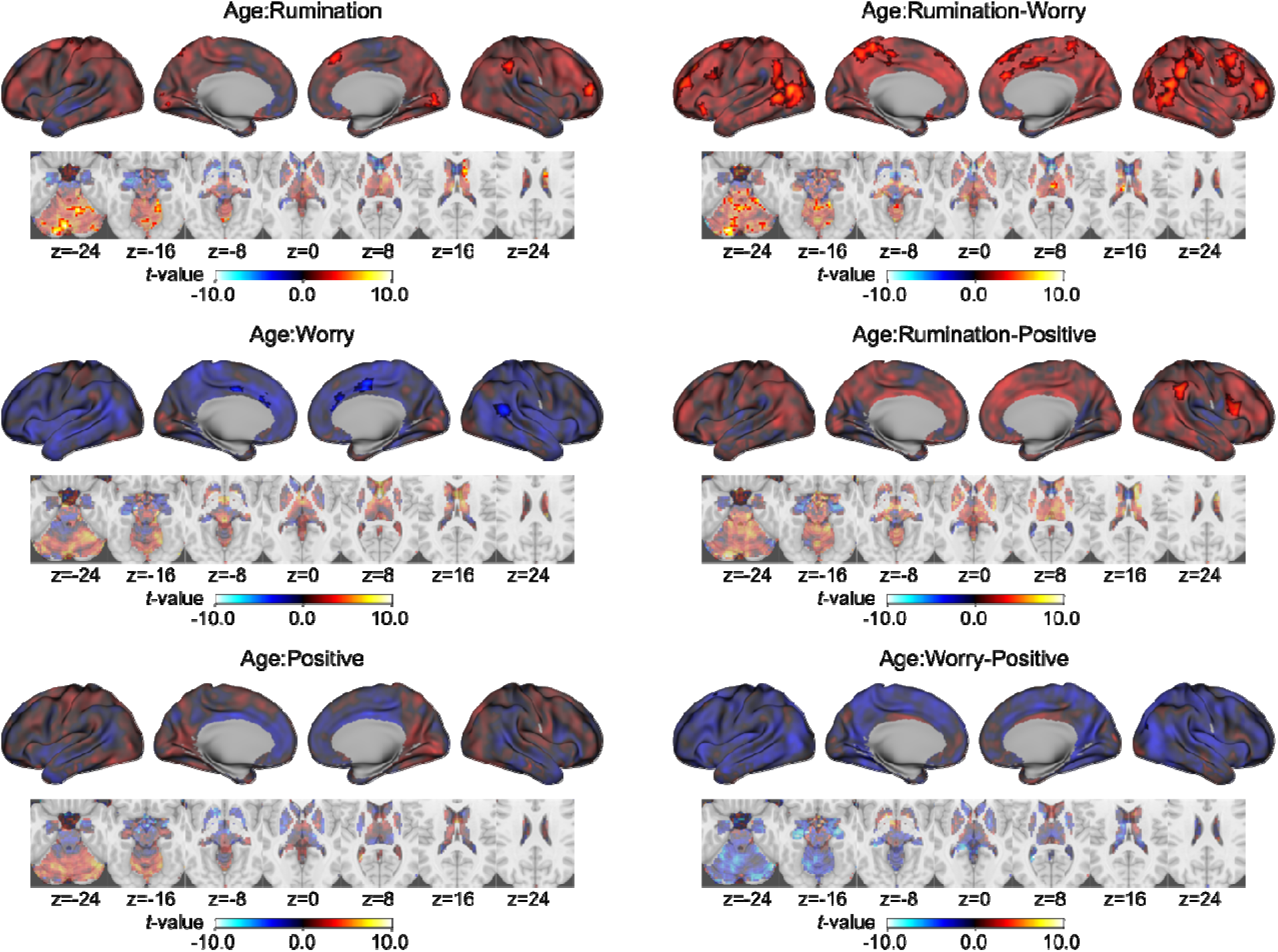
Brain activation patterns associated with age in each thought state and their contrasts. The maps are displayed on the inflated cortical surface, with subcortical and cerebellar regions shown on axial slice maps. Clusters with voxel-wise p < 0.001, corrected for cluster extent at p < 0.05, are highlighted with opaque colors. Unthresholded maps are presented with lower opacity in the plots. The images are displayed in neurological orientation, with the left side corresponding to the left side of the brain.

In contrast to the findings on rumination, there was a decline in brain activity associated with worry as individuals aged, particularly in the right TPJ and anterior and middle cingulate regions. The TPJ is a crucial brain area involved in a range of cognitive functions, particularly those related to social cognition. Specifically, the right TPJ plays a pivotal role in understanding others’ mental states ^34–36^. Its role in processing self-referential and social information may contribute to the cognitive aspects of worry and anxiety, as these states often involve heightened self-focus and concerns about social evaluation. The TPJ’s connectivity with regions involved in emotional processing, such as the anterior insula and the prefrontal cortex, provides further evidence of its involvement in these affective states ^36,37^. Increased activity in the anterior and middle cingulate regions has previously been linked to anxiety ^38,39^ and social distress ^40^.

These findings suggest that, with age, there is a reduction in brain activity associated with future-oriented anxiety about social relationships. Given that RNT often pertains to social relationships and self-evaluation ^20^, the observed decline in these brain activations may be associated with a reduction in anxiety responses to social events. This is also in line with the fact that older adults have lower levels of affective reactivity than younger adults ^3^. Furthermore, research indicates that emotional empathy tends to increase with age ^41,42^, suggesting that the reduction in social anxiety in older adults may be an adaptive change.

Changes in emotional processing with age may not reflect cognitive decline but rather adaptive shifts in the functional significance of emotions ^43^ and age-related goals influenced by changes in the social environment. Socioemotional Selectivity Theory (SST) suggests that as we age, we prioritize emotionally meaningful goals, savoring positive emotions while avoiding negative ones ^44^. SST posits that as individuals perceive the end of life approaching, emotionally meaningful goals take precedence over exploration. The present findings complement this theory by demonstrating that the age-related decline in negative thinking is driven by distinct neural changes in rumination and worry. These results highlight the importance of examining emotional aging not only through the general valence of emotions but also by considering how aging affects individual emotions as discrete processes ^43^. This perspective offers a more nuanced understanding of emotional aging, suggesting that the decline in negative thinking is tied to specific adaptive processes for each emotional state, rather than a simple shift toward positive or negative emotions.

Although the present results were based on healthy subjects, age influences the neural substrates relevant to depression, with previous research indicating distinct variations across age groups in major depressive disorder (MDD). Specifically, alterations in the global network organization of structural and functional brain connectomes in MDD vary according to the age of onset ^45^. Late-onset depression, in particular, is more strongly associated with anatomical brain alterations compared to early-onset depression ^46–48^. The present findings demonstrate that the RNT process also exhibits age-related variations, with distinct effects on rumination and worry. Research suggests that these two processes differentially impact mood disorders: rumination is associated with depressive comorbidity and recurrence, while worry is more closely linked to anxiety disorders ^17,18^. Therefore, age-related differences may also influence the relationship between RNT and depression. Future studies should explore how age interacts with these processes to better understand their role in mood disorders.

It is important to highlight the methodological strengths of this study. We analyzed brain activity using decoding probability outputs at each time point as regressors. The observed average activity patterns (Figure 5) closely matched those from traditional block-model analyses (Supplementary Figure S1), validating our approach. Moreover, utilizing decoding probability allowed us to capture dynamic changes in brain activity, offering more precise and individualized modeling of emotional state transitions compared to static, predefined state models. The effectiveness of machine learning in detecting and tracking changes in mental states over time has been demonstrated in other studies ^49,50^. In the present study, this approach successfully highlighted individual differences in the time course of decoded states and revealed variability within individuals across sessions and trials. By accounting for these variations, our regression analysis provided a more nuanced detection of brain activity, likely contributing to the identification of age-related effects in this study.

This study has several limitations. First, the small sample size limits the generalizability of the findings. A larger sample would provide greater statistical power and a more robust examination of the effects of age on emotional brain states. Second, the participant pool was female-biased, which may influence the results, as sex differences have been observed in emotional processing ^51^. Future studies should aim to include a more balanced sample to better understand the role of gender in these processes. Finally, this study was cross-sectional, making it difficult to draw conclusions about the temporal dynamics of age-related changes in emotional processing. Longitudinal studies are needed to track how rumination, worry, and their neural correlates evolve over time and to determine whether the observed changes are causally linked to aging.

In conclusion, the present results suggest that emotional brain states evolve across the lifespan, enhancing emotional well-being in older adults, particularly through changes in negative thinking processes. Notably, these changes are not uniform across individual negative emotional states such as rumination and worry. Aging appears to enhance cognitive control during rumination and reduce neural responses to anxiety-provoking events during worry. The individuality of emotional brain states and their age-related evolution should be a key consideration in promoting lifelong mental health and developing personalized interventions for mental disorders. Understanding these dynamics could pave the way for more targeted and effective mental health strategies across different life stages.

## Methods

### Participants

Thirty-seven healthy individuals (24 females, aged 18–64) participated in the study.

Exclusion criteria included a positive to drugs abuse test, a lifetime diagnosis of psychiatric disorders, and significant issues that could affect participation, such as uncorrectable vision or hearing problems and MRI contraindications. Individuals with moderate to severe traumatic brain injury, active suicidal ideation, recent changes in medication affecting brain function, or prescriptions outside of accepted ranges were also excluded. All participants provided informed consent in accordance with the principles outlined in the Declaration of Helsinki. The study procedures were approved by the WCG IRB (https://www.wcgclinical.com; tracking number: 20224917).

### Session schedule and assessment measures

The study sessions consisted of three visits: one preparation session and two MRI scanning sessions. During the first visit, participants received study information, provided informed consent, and completed assessments for negative and positive thinking tendencies. These included the Ruminative Response Scale (RRS) ^28^, the Brief State Rumination Inventory (BSRI) ^31^, and the State-Trait Anxiety Inventory (STAI) ^29^. Positive affect was measured using the Spinal Cord Injury Quality of Life (SCI-QOL) Positive Affect and Well-Being score ^30^, part of the NIH Toolbox for Emotion Assessment ^52^. State scores (BSRI, STAI-S) were also collected before the fMRI session on the second visit.

Although participants were healthy, assessments for depression and anxiety were conducted using the Quick Inventory of Depressive Symptomatology (QIDS) ^53^ and the Penn State Worry Questionnaire (PSWQ) ^54^. Additionally, emotion regulation abilities were measured using the Metacognitive Questionnaire-30 (MCQ-30) ^55^, the Thought Control Questionnaire (TCQ) ^56^, the Emotion Regulation Questionnaire (ERQ), the Difficulties in Emotion Regulation Scale (DERS) ^57^, and the Cognitive Emotion Regulation Questionnaire (CERQ) ^58^, aiming to associate these measures with performance in an emotion regulation task conducted during a third visit, which is not included in the present paper. The primary focus of this study was on the RRS, BSRI, STAI, and SCI-QOL Positive measures to analyze the relationship between brain state discriminability for rumination, worry, and positive thinking and their associated psychological measures. Other measures were analyzed as exploratory variables (see Supplementary Table S2).

During the second visit, participants were asked to recall twelve autobiographical events, each clearly associated with one of three different thinking patterns: rumination ("thoughts you have had about the past that make you feel bad, embarrassed, sad, or afraid when you think about them, and that may be hard to stop thinking about"), worry ("thoughts that may make you anxious, nervous, tense, or irritable, and sometimes it is difficult to get these thoughts out of your mind"), and positive thinking ("thoughts you have had about the past that make you feel good, that you enjoy thinking about, and that are happy, positive, fulfilling, or exciting events in your life"). Participants generated four events for each category and wrote down short descriptions (keywords) to help them recall these events during the MRI session. To maintain confidentiality, no details of the personal events were requested, allowing participants to honestly recall their negative personal experiences. Only distress levels associated with the rumination and worry events were collected using a seven-point Likert scale (1: Not distressing at all, to 7: Extremely distressing).

Participants also completed the state rumination scale (BSRI) ^31^ and the state anxiety scale (STAI-S) ^29^ prior to the MRI scanning session and these scores were used in the analysis (Table 2 and Figure 3). During the MRI session, participants performed the Thought Induction Task (TIT) with functional MRI (fMRI), as detailed below. In the third visit, participants completed another emotion regulation task, which is not included in the present paper and will be reported separately.

### MRI scanning parameters

The MRI scan was performed using a 3T Discovery MR750 scanner (GE Healthcare, Milwaukee, WI, USA) with a 32-channel head coil. Brain functional images of blood oxygenation level-dependent (BOLD) signals were collected using gradient echo-planar imaging of T2*- weighted signals with the following parameters: TR = 2.0 s, TE = 25 ms, FA = 90°, SENSE acceleration factor (R) = 2, number of axial slices = 40, slice thickness = 2.9 mm, FOV = 240 × 240 mm, and matrix = 96 × 96, resampled to 128 × 128.

The anatomical reference image of the brain was obtained using T1-weighted imaging with the MPRAGE sequence with TR = 6 ms, TE = 2.92 ms, SENSE acceleration factor (R) = 2, flip angle = 8°, inversion time = 1060 ms, sampling bandwidth = 31.25 kHz, FOV = 256 × 256 mm, 208 sagittal slices, slice thickness = 1.0 mm, and a scan time of 6 min 11 s.

### Thought Induction Task (TIT)

After preparation runs for locating the imaging position, the scan session began with the T1- weighted anatomical scan, followed by the resting-state run (6 min 50 s), and then six runs of the Thought Induction Task (TIT), each lasting 8 min 2 s (Figure 1). Each run of the TIT began with an 18-s Rest block, followed by a 22-s Think block and an 18-s Flanker task block. These blocks were repeated eight times per run and concluded with an 18-s Rest block.

During the Rest block, a white cross was displayed on the screen, and participants were instructed not to think about anything in particular. During the Think block, keywords representing one of the personal events collected prior to the scan were shown along with an instruction sentence, such as "Why did that happen to you?" or "Why did you do that at the moment?" for rumination. The instruction sentence appropriate for each event was selected by the participants during the thought collection process. Participants were instructed to engage with the thought related to the event’s keywords according to the instruction sentence.

In the Flanker task block, five arrowheads were presented, and participants were instructed to respond to the direction of the center arrow by pressing the left or right button as quickly as possible. The four flanker arrows pointed in the same direction, while the center arrow could point in the same (congruent) or opposite (incongruent) direction. Once the button was pressed, the arrows disappeared without any feedback. Trials were presented every 3 seconds, regardless of response time, and six trials were performed in each block. This task was designed to impose an attentional load and was placed after the Think block to help participants clear their thoughts before the next Rest block.

Six TIT runs were performed during the session. Each of the 12 thought events was presented four times during the session, and the order of events was pseudorandomized with the following restrictions: events from the same category (rumination, worry, positive) did not repeat consecutively, each run included approximately the same number of events (two to three) from each category, and the same event did not repeat within the same category.

At the end of each task run, participants answered five questions about their mental states during and after the task using a seven-point Likert scale: ’How difficult was it to engage with your thoughts during the Think blocks?’ (difficulty), ’How successful were you in engaging with your thoughts during the Think blocks?’ (success), ’How sleepy were you during the scan?’ (sleepiness), ’How emotionally drained do you feel right now?’ (emotional exhaustion), and ’How tired are you right now?’ (tiredness).

### MRI data preprocessing

We used AFNI (https://afni.nimh.nih.gov/) to process the MRI image data. The initial three fMRI volumes were discarded to allow the signal to reach a steady state. The following preprocessing steps were applied: despiking, RETROICOR ^59^ and RVT ^60^ physiological noise correction, slice-timing correction, motion correction, spatial normalization to the MNI template brain with resampling to a 3 × 3 × 3 mm voxel size using Advanced Normalization Tools (ANTs) ^61^, spatial smoothing with a 6 mm full-width at half maximum (FWHM) kernel within the brain, and scaling to percent signal change in each voxel.

Next, noise components were regressed out using general linear model (GLM) analysis.

The noise components included low-frequency drift modeled by up to third-order Legendre polynomials, six motion parameters (three rotations and three translations) and their temporal derivatives, three principal components of the ventricle signals, and local white matter average signals (ANATICOR) ^62^. Volumes with frame-wise displacement greater than 0.3 mm and their preceding volumes were censored in the GLM analysis. Task runs with more than one-third of time points were censored were excluded from the analysis. Response models for each of the Flanker task trials were also included, with congruent and incongruent trials modeled separately using a canonical hemodynamic response function (HRF) convolved with a delta function at the onset of each event. The residuals from the GLM analysis were used as input for the decoding analysis of thought states.

### Decoding thought states

The thought state at each time point during the Think and Rest blocks were classified using a machine learning model. The input signals were extracted from the gray matter voxels of the preprocessed images across the whole brain. The data for each thought state were sampled during the Think block, excluding the initial 3 time points (6 s) to account for the hemodynamic response delay, providing 8 time points per block and 128 samples per thought category. Data for the rest state were sampled during the Rest block, excluding the initial 6 time points (12 s) to avoid any residual effects from the Flanker task and to approximately equalize the number of samples with the other thought states. This provided 3 time points per block and 144 samples in total. Censored time points were excluded from the input to the decoding analysis.

The classification analysis was performed using the Auto-sklearn package (https://automl.github.io/auto-sklearn/) ^27^. This package runs an AutoML analysis, which performs Combined Algorithm Selection and Hyperparameter optimization (CASH) to find the optimal model. Auto-sklearn creates ensembles of data preprocessors, feature preprocessors, and machine learning models from the scikit-learn (https://scikit-learn.org) machine learning package in Python. The model training was performed using default parameters, except for the time limit for model training. In the default configuration, the Auto-sklearn search was time-limited to 6 minutes per model training call and 1 hour for the entire task, given the extensive parameter space in the CASH process. We extended these limits to 3 hours per model training call and 30 hours for the total task to increase the likelihood of identifying better-performing models. Additionally, we tested limits of 6 hours per model call and 60 hours in total, but observed no significant performance improvement, suggesting that model performance plateaued with the 3-hour and 30-hour configuration.

Training and evaluation were performed using leave-one-run-out cross-validation, where a model was trained on samples from five TIT runs and tested on the remaining run. This process was repeated for each of the six runs, with each run used as the test set once. The model was optimized using nested leave-one-run-out cross-validation within the training samples, while the test sample was reserved exclusively for evaluating model performance. Model performance was assessed using the area under the receiver operating characteristic curve (AUC).

Classification accuracies were not used, as they could be biased by sample size imbalances. Since motion censoring could lead to such imbalances between thought categories, classification accuracies are difficult to compare across participants.

The time course of the model outputs across all time points in a test run was used to infer the series of thought states across blocks. We considered this time series to be more sensitive in tracking participants’ thought states, as it could represent individualized and dynamic mental state evolution ^50^, compared to a block-wise model, which assumes a stable state during the Think block. Therefore, this time series of decoded thought states was used as GLM regressors to evaluate brain activation patterns associated with each decoded thought state. This GLM analysis included all previously mentioned noise components and was applied to the preprocessed fMRI data. As an exploratory analysis, we also ran a GLM with the block response model (a boxcar function convolved with HRF) for the Think blocks of each category to compare the results between the decoded time course and the block regressors.

### Statistical analysis

The significance of AUC for thought state discrimination was assessed for each participant using a normal approximation of Mann-Whitney U statistics ^63^. The associations between thought state discriminability (AUC) and each of the demographic variables and assessment scores were tested using the Spearman correlation, except for sex, where a t-test was used.

The *p*-values across these variables were corrected by false discovery rate (FDR) ^64^.

The effect of age group on the temporal pattern of decoding state probability was tested using linear mixed-effects (LME) model analysis with the state probability of the corresponding thought as the dependent variable and the fixed effects of time, age group (divided by median age = 39), and their interaction, and a random intercept for participants. For this analysis, the time points were restricted to the samples used for model training (from 6 seconds after block onset to the end of the block).

Group analysis of the BOLD signal change associated with thought states was performed using LME model analysis, with the beta maps of each thought condition for each run, obtained from the first-level GLM analysis, as the dependent variables, the thought condition, participants’ age, and their interaction as the independent variables, and participant as a random effect on the intercept. The LME analysis was performed using the *lme4* package ^65^ in the R language for statistical computing, and contrast maps were computed using the *emmeans* package ^66^. The statistical map from the LME analysis was thresholded at *p* < 0.001 voxel-wise, and a cluster extent threshold of *p* < 0.05 was then applied. The cluster extent threshold was evaluated using AFNI 3dClustSim with an improved spatial autocorrelation function ^67^.

