## Supplementary Information for "Aging increases the distinctiveness of emotional brain states across rumination, worry, and positive thinking"

**Supplementary Table S1.** AUC correlations with self-reports of task evaluation and distress levels related to rumination and worry events. P-values are uncorrected.

| | Rumination-Worry | | Rumination-Positive | | Worry-Positive | |
|---|---|---|---|---|---|---|
| | Spearman rho | *p* | Spearman rho | *p* | Spearman rho | *p* |
| difficulty | -0.111 | 0.525 | -0.190 | 0.273 | -0.291 | 0.089 |
| success | 0.054 | 0.756 | 0.174 | 0.318 | 0.254 | 0.141 |
| sleepiness | -0.151 | 0.386 | -0.273 | 0.113 | -0.275 | 0.110 |
| emotional exhaustion | 0.099 | 0.573 | 0.050 | 0.776 | 0.076 | 0.665 |
| tiredness | -0.178 | 0.306 | -0.262 | 0.128 | -0.210 | 0.225 |
| distress rumination | 0.034 | 0.848 | 0.168 | 0.336 | 0.229 | 0.186 |
| distress worry | 0.280 | 0.103 | 0.238 | 0.168 | 0.184 | 0.290 |

**Supplementary Table S2**. AUC correlations with exploratory measures. P-values are uncorrected.

| | Rumination-Worry | | Rumination-Positive | | Worry-Positive | |
|---|---|---|---|---|---|---|
| | Spearman rho | *p* | Spearman rho | *p* | Spearman rho | *p* |
| QIDS-SR | -0.143 | 0.412 | -0.290 | 0.091 | -0.208 | 0.231 |
| PSWQ | -0.045 | 0.799 | -0.208 | 0.231 | -0.104 | 0.553 |
| MCQ-30 | -0.035 | 0.841 | -0.300 | 0.080 | -0.271 | 0.115 |
| TCQ | 0.127 | 0.468 | -0.081 | 0.644 | -0.012 | 0.946 |
| ERQ | 0.202 | 0.244 | 0.108 | 0.537 | 0.183 | 0.294 |
| DERS | -0.332 | 0.052 | -0.490 | 0.003* | -0.399 | 0.017* |
| CERQ | 0.129 | 0.462 | -0.034 | 0.847 | 0.041 | 0.814 |

*, $p$ (uncorrected) < 0.05; QIDS, Quick Inventory of Depressive Symptomatology via self-assessment; PSWQ, Penn State Worry Questionnaire; MCQ-30, Metacognitive Questionaire-30; TCQ, Thought Control Questionnaire, ERQ, Emotion Regulation Questionnaire; DERS, Difficulties in Emotion Regulation Scale; CERQ, Cognitive Emotion Regulation Questionnaire.

**Supplementary Table S3.** Peak coordinates of significant clusters of brain activation during the rumination state. Peak points at least 12 mm apart were extracted from each cluster.

| Cluster ID | X | Y | Z | Peak Stat | Cluster Size ($mm^3$) | Area |
|---|---|---|---|---|---|---|
| | | | | | Positive clusters | |
| 1 | -4.5 | 7.5 | 61.5 | 11.128 | 34695 | Left SMA |
| 1a | -19.5 | 13.5 | 7.5 | 7.738 | | Left Putamen |
| 1b | -13.5 | 19.5 | 37.5 | 7.398 | | Left Middle Cingulate Cortex |
| 1c | -7.5 | 52.5 | 34.5 | 7.263 | | Left Superior Medial Gyrus |
| 2 | -16.5 | -91.5 | -1.5 | 10.026 | 14553 | Left Middle Occipital Gyrus |
| 2a | -40.5 | -73.5 | -13.5 | 8.425 | | Left Inferior Occipital Gyrus |
| 2b | -28.5 | -82.5 | -10.5 | 8.028 | | Left Inferior Occipital Gyrus |
| 2c | -37.5 | -46.5 | -19.5 | 8.006 | | Left Fusiform Gyrus |

| | | | | | | |
|---|---|---|---|---|---|---|
| 3 | 16.5 | -91.5 | -1.5 | 9.909 | 14796 | Right Calcarine Gyrus |
| 3a | 25.5 | -79.5 | -7.5 | 7.761 | | Right Lingual Gyrus |
| 3b | 31.5 | -67.5 | -22.5 | 7.381 | | Right Cerebellum (VI) |
| 3c | 40.5 | -73.5 | -13.5 | 5.233 | | Right Inferior Occipital Gyrus |
| 4 | -37.5 | 28.5 | -1.5 | 9.698 | 66798 | Left Inferior Frontal Gyrus (p. Triangularis) |
| 4a | -46.5 | 1.5 | 52.5 | 9.648 | | Left Precentral Gyrus |
| 4b | -37.5 | 34.5 | -1.5 | 9.484 | | Left Inferior Frontal Gyrus (p. Triangularis) |
| 4c | -49.5 | 22.5 | -1.5 | 9.185 | | Left Inferior Frontal Gyrus (p. Triangularis) |
| 5 | -28.5 | -25.5 | -4.5 | 6.917 | 1242 | Left Hippocampus |
| 6 | 43.5 | -31.5 | 4.5 | 6.706 | 6966 | Right Superior Temporal Gyrus |
| 6a | 46.5 | -16.5 | -10.5 | 6.228 | | Right Superior Temporal Gyrus |
| 6b | 49.5 | 16.5 | -16.5 | 5.677 | | Right Temporal Pole |
| 6c | 46.5 | -13.5 | -16.5 | 5.209 | | Right Middle Temporal Gyrus |
| 7 | 58.5 | 7.5 | 37.5 | 6.508 | 4131 | Right Precentral Gyrus |
| 7a | 52.5 | 1.5 | 52.5 | 6.246 | | Right Middle Frontal Gyrus |
| 7b | 34.5 | 1.5 | 40.5 | 4.164 | | Right Middle Frontal Gyrus |
| 8 | 43.5 | 28.5 | 1.5 | 6.228 | 5535 | Right Inferior Frontal Gyrus (p. Triangularis) |
| 8a | 28.5 | 34.5 | -1.5 | 5.466 | | Right Inferior Frontal Gyrus (p. Orbitalis) |
| 8b | 28.5 | 22.5 | -13.5 | 4.753 | | Right Insula Lobe |
| 8c | 58.5 | 25.5 | 22.5 | 4.715 | | Right Inferior Frontal Gyrus (p. Triangularis) |
| 9 | 19.5 | 10.5 | 7.5 | 5.615 | 1404 | Right Putamen |
| 9a | 19.5 | 4.5 | 19.5 | 4.630 | | Right Caudate Nucleus |
| 10 | -7.5 | -22.5 | -4.5 | 4.880 | 756 | Left Thalamus |
| Negative clusters | | | | | | |
| 1 | 58.5 | -46.5 | 43.5 | -8.740 | 14715 | Right Inferior Parietal Lobule |
| 1a | 43.5 | -76.5 | 37.5 | -6.116 | | Right Angular Gyrus |
| 1b | 61.5 | -31.5 | 49.5 | -5.780 | | Right SupraMarginal Gyrus |
| 1c | 43.5 | -67.5 | 52.5 | -5.719 | | Right Angular Gyrus |
| 2 | 1.5 | -37.5 | 49.5 | -8.734 | 10044 | Left Middle Cingulate Cortex |
| 2a | -10.5 | -34.5 | 43.5 | -6.626 | | Left Middle Cingulate Cortex |
| 3 | -43.5 | -7.5 | -1.5 | -7.733 | 3753 | Left Insula Lobe |
| 3a | -37.5 | 7.5 | -10.5 | -4.218 | | Left Insula Lobe |
| 4 | 31.5 | 28.5 | 43.5 | -7.098 | 7668 | Right Middle Frontal Gyrus |
| 4a | 25.5 | 16.5 | 61.5 | -6.351 | | Right Superior Frontal Gyrus |
| 4b | 19.5 | 22.5 | 49.5 | -5.835 | | Right Superior Frontal Gyrus |
| 5 | 43.5 | 49.5 | -1.5 | -7.047 | 7776 | Right Middle Orbital Gyrus |
| 5a | 43.5 | 40.5 | 13.5 | -6.654 | | Right Middle Frontal Gyrus |
| 5b | 40.5 | 55.5 | -1.5 | -6.527 | | Right Middle Orbital Gyrus |
| 5c | 22.5 | 67.5 | 1.5 | -5.371 | | Right Superior Frontal Gyrus |
| 6 | 4.5 | 49.5 | -4.5 | -5.986 | 6453 | Right Mid Orbital Gyrus |

| | | | | | | |
|---|---|---|---|---|---|---|
| 6a | 7.5 | 37.5 | 10.5 | -5.806 | | Right Anterior Cingulate Cortex |
| 6b | 16.5 | 34.5 | -13.5 | -5.283 | | Right Superior Orbital Gyrus |
| 6c | 13.5 | 40.5 | 13.5 | -4.787 | | Right Anterior Cingulate Cortex |
| 7 | -67.5 | -31.5 | 40.5 | -5.874 | 1782 | Left SupraMarginal Gyrus |
| 7a | -67.5 | -31.5 | 25.5 | -4.692 | | Left SupraMarginal Gyrus |
| 8 | -34.5 | -43.5 | -7.5 | -5.369 | 918 | Left ParaHippocampal Gyrus |
| 9 | 55.5 | -61.5 | 1.5 | -4.904 | 1539 | Right Middle Temporal Gyrus |
| 9a | 58.5 | -52.5 | -7.5 | -4.637 | | Right Inferior Temporal Gyrus |
| 10 | -16.5 | -34.5 | -13.5 | -4.493 | 675 | Left ParaHippocampal Gyrus |
| 10a | -22.5 | -46.5 | -10.5 | -4.415 | | Left Fusiform Gyrus |

**Supplementary Table S4.** Peak coordinates of significant clusters of brain activation during the worry state. Peak points at least 12 mm apart were extracted from each cluster.

| Cluster ID | X | Y | Z | Peak Stat | Cluster Size (mm$^3$) | Area |
|---|---|---|---|---|---|---|
| | | | | | Positive clusters | |
| 1 | -16.5 | -91.5 | -1.5 | 12.831 | 64665 | Left Middle Occipital Gyrus |
| 1a | 16.5 | -91.5 | -1.5 | 11.669 | | Right Calcarine Gyrus |
| 1b | -40.5 | -73.5 | -13.5 | 10.990 | | Left Inferior Occipital Gyrus |
| 1c | -19.5 | -85.5 | -7.5 | 10.719 | | Left Fusiform Gyrus |
| 2 | -7.5 | 10.5 | 64.5 | 11.834 | 37881 | Left SMA |
| 2a | -10.5 | 22.5 | 64.5 | 10.722 | | Left SMA |
| 2b | -10.5 | 46.5 | 43.5 | 9.748 | | Left Superior Frontal Gyrus |
| 2c | -7.5 | 37.5 | 52.5 | 6.684 | | Left Superior Medial Gyrus |
| 3 | -46.5 | 1.5 | 52.5 | 10.777 | 56592 | Left Precentral Gyrus |
| 3a | -52.5 | 19.5 | 19.5 | 10.348 | | Left Inferior Frontal Gyrus (p. Triangularis) |
| 3b | -40.5 | 28.5 | -4.5 | 9.854 | | Left Inferior Frontal Gyrus (p. Orbitalis) |
| 3c | -58.5 | 22.5 | 10.5 | 9.759 | | Left Inferior Frontal Gyrus (p. Triangularis) |
| 4 | -25.5 | -28.5 | -1.5 | 10.266 | 2889 | Left Hippocampus |
| 5 | 25.5 | -25.5 | -1.5 | 9.477 | 1350 | Right Thalamus |
| 6 | -55.5 | -37.5 | 1.5 | 9.030 | 12636 | Left Middle Temporal Gyrus |
| 6a | -46.5 | -28.5 | -4.5 | 8.596 | | Left Middle Temporal Gyrus |
| 6b | -67.5 | -40.5 | 1.5 | 7.168 | | Left Middle Temporal Gyrus |
| 7 | -19.5 | 13.5 | 7.5 | 7.702 | 11691 | Left Putamen |
| 7a | -16.5 | -10.5 | 25.5 | 5.718 | | Left Caudate Nucleus |
| 7b | -10.5 | -13.5 | 4.5 | 4.890 | | Left Thalamus |
| 7c | -7.5 | -22.5 | -10.5 | 4.556 | | Left ParaHippocampal Gyrus |
| 8 | 55.5 | 25.5 | 7.5 | 7.248 | 7641 | Right Inferior Frontal Gyrus (p. Triangularis) |
| 8a | 49.5 | 25.5 | -10.5 | 5.744 | | Right Inferior Frontal Gyrus (p. Orbitalis) |
| 8b | 43.5 | 10.5 | 4.5 | 4.462 | | Right Insula Lobe |

| | | | | | | |
|---|---|---|---|---|---|---|
| 8c | 34.5 | 25.5 | -7.5 | 4.360 | | Right Inferior Frontal Gyrus (p. Orbitalis) |
| 9 | 43.5 | -34.5 | 4.5 | 6.747 | 3402 | Right Superior Temporal Gyrus |
| 9a | 46.5 | -19.5 | -7.5 | 4.207 | | Right Middle Temporal Gyrus |
| 10 | 55.5 | -1.5 | 49.5 | 6.401 | 6885 | Right Precentral Gyrus |
| 10a | 64.5 | -1.5 | 28.5 | 5.683 | | Right Postcentral Gyrus |
| 11 | -46.5 | -58.5 | 31.5 | 5.821 | 3699 | Left Angular Gyrus |
| 11a | -58.5 | -49.5 | 28.5 | 5.292 | | Left SupraMarginal Gyrus |
| 12 | 40.5 | 4.5 | -34.5 | 5.747 | 3753 | Right Medial Temporal Pole |
| 12a | 52.5 | 13.5 | -28.5 | 5.478 | | Right Medial Temporal Pole |
| 12b | 31.5 | 16.5 | -31.5 | 4.697 | | Right Temporal Pole |
| 12c | 49.5 | 16.5 | -19.5 | 4.587 | | Right Temporal Pole |
| 13 | -1.5 | 55.5 | -19.5 | 5.340 | 1323 | Left Rectal Gyrus |
| 14 | 22.5 | 4.5 | 10.5 | 5.232 | 2538 | Right Putamen |
| 14a | 19.5 | 16.5 | 7.5 | 5.000 | | Right Caudate Nucleus |
| 14b | 16.5 | -1.5 | 1.5 | 4.987 | | Right Pallidum |
| 14c | 22.5 | -1.5 | 13.5 | 4.898 | | Right Putamen |
| 15 | 19.5 | 52.5 | 40.5 | 5.167 | 2835 | Right Superior Frontal Gyrus |
| 15a | 7.5 | 61.5 | 34.5 | 4.818 | | Right Superior Medial Gyrus |
| 15b | 16.5 | 49.5 | 28.5 | 4.506 | | Right Superior Frontal Gyrus |
| 16 | -7.5 | -49.5 | 25.5 | 4.994 | 1350 | Left Posterior Cingulate Cortex |
| | | | | Negative clusters | | |
| 1 | -43.5 | -10.5 | -1.5 | -9.340 | 6534 | Left Superior Temporal Gyrus |
| 1a | -46.5 | 4.5 | -10.5 | -5.268 | | Left Superior Temporal Gyrus |
| 1b | -52.5 | -1.5 | -1.5 | -5.208 | | Left Superior Temporal Gyrus |
| 1c | -40.5 | -1.5 | -10.5 | -4.986 | | Left Superior Temporal Gyrus |
| 2 | -13.5 | -37.5 | 43.5 | -8.059 | 14283 | Left Middle Cingulate Cortex |
| 2a | 4.5 | -37.5 | 46.5 | -7.436 | | Right Middle Cingulate Cortex |
| 2b | 10.5 | -58.5 | 52.5 | -5.580 | | Right Precuneus |
| 2c | -10.5 | -52.5 | 58.5 | -4.956 | | Left Precuneus |
| 3 | 61.5 | -28.5 | 40.5 | -6.651 | 3780 | Right SupraMarginal Gyrus |
| 3a | 61.5 | -43.5 | 43.5 | -4.606 | | Right SupraMarginal Gyrus |
| 4 | -58.5 | -31.5 | 34.5 | -6.645 | 2349 | Left SupraMarginal Gyrus |
| 4a | -64.5 | -28.5 | 46.5 | -5.761 | | Left SupraMarginal Gyrus |
| 5 | 46.5 | -73.5 | 25.5 | -6.296 | 2160 | Right Middle Occipital Gyrus |
| 5a | 37.5 | -79.5 | 43.5 | -5.891 | | Right Superior Occipital Gyrus |
| 6 | -31.5 | -85.5 | 37.5 | -5.896 | 1188 | Left Middle Occipital Gyrus |
| 6a | -43.5 | -82.5 | 31.5 | -5.094 | | Left Middle Occipital Gyrus |
| 7 | -37.5 | -43.5 | -7.5 | -5.770 | 1944 | Left Inferior Temporal Gyrus |
| 7a | -13.5 | -37.5 | -10.5 | -5.130 | | Left Fusiform Gyrus |
| 7b | -28.5 | -43.5 | -10.5 | -4.824 | | Left Fusiform Gyrus |

| | | | | | | |
|---|---|---|---|---|---|---|
| 7c | -13.5 | -31.5 | -10.5 | -4.524 | | Left ParaHippocampal Gyrus |
| 8 | 16.5 | -34.5 | -10.5 | -5.729 | 675 | Right Lingual Gyrus |
| 8a | 7.5 | -37.5 | -19.5 | -4.452 | | Cerebellar Vermis (1/2) |
| 9 | 49.5 | -52.5 | -1.5 | -5.541 | 1107 | Right Middle Temporal Gyrus |
| 10 | 22.5 | -52.5 | 19.5 | -5.335 | 891 | Right Precuneus |
| 11 | 40.5 | -7.5 | -1.5 | -5.232 | 1404 | Right Insula Lobe |
| 11a | 40.5 | -10.5 | -13.5 | -5.133 | | Right Hippocampus |

**Supplementary Table S5.** Peak coordinates of significant clusters of brain activation during the positive thinking state. Peak points at least 12 mm apart were extracted from each cluster.

| Cluster ID | X | Y | Z | Peak Stat | Cluster Size (mm$^3$) | Area |
|---|---|---|---|---|---|---|
| | | | | Positive clusters | | |
| 1 | -16.5 | -91.5 | -1.5 | 11.618 | 131031 | Left Middle Occipital Gyrus |
| 1a | 16.5 | -91.5 | -1.5 | 11.543 | | Right Calcarine Gyrus |
| 1b | -25.5 | -28.5 | -1.5 | 10.555 | | Left Hippocampus |
| 1c | -46.5 | 16.5 | -25.5 | 9.845 | | Left Medial Temporal Pole |
| 2 | -4.5 | 7.5 | 61.5 | 9.866 | 12879 | Left SMA |
| 3 | 25.5 | -25.5 | -1.5 | 9.264 | 1296 | Right Thalamus |
| 4 | 31.5 | 10.5 | -34.5 | 7.253 | 14796 | Right Medial Temporal Pole |
| 4a | 43.5 | 19.5 | -31.5 | 6.758 | | Right Temporal Pole |
| 4b | 58.5 | 4.5 | -19.5 | 6.675 | | Right Middle Temporal Gyrus |
| 4c | 43.5 | 16.5 | -37.5 | 6.490 | | Right Medial Temporal Pole |
| 5 | 31.5 | 34.5 | -10.5 | 6.914 | 1620 | Right Inferior Frontal Gyrus (p. Orbitalis) |
| 6 | -7.5 | 55.5 | -10.5 | 6.396 | 3645 | Left Mid Orbital Gyrus |
| 6a | 7.5 | 55.5 | -10.5 | 4.968 | | Right Mid Orbital Gyrus |
| 6b | -7.5 | 64.5 | 7.5 | 4.866 | | Left Superior Medial Gyrus |
| 7 | -10.5 | 52.5 | 37.5 | 6.016 | 2187 | Left Superior Frontal Gyrus |
| 8 | 55.5 | 1.5 | 49.5 | 5.364 | 4995 | Right Precentral Gyrus |
| 8a | 61.5 | -1.5 | 34.5 | 5.260 | | Right Postcentral Gyrus |
| 8b | 46.5 | -7.5 | 34.5 | 4.990 | | Right Precentral Gyrus |
| 8c | 64.5 | 4.5 | 22.5 | 4.352 | | Right Precentral Gyrus |
| 9 | 4.5 | 16.5 | -16.5 | 5.354 | 1512 | Right Olfactory cortex |
| 9a | 1.5 | 4.5 | -16.5 | 5.067 | | Left Olfactory cortex |
| 9b | -1.5 | 16.5 | -7.5 | 4.826 | | Left Olfactory cortex |
| 10 | -58.5 | -43.5 | -7.5 | 5.197 | 2565 | Left Middle Temporal Gyrus |
| 10a | -49.5 | -40.5 | 7.5 | 4.532 | | Left Middle Temporal Gyrus |
| 11 | -28.5 | -73.5 | 49.5 | 4.583 | 891 | Left Inferior Parietal Lobule |
| 11a | -31.5 | -67.5 | 34.5 | 3.828 | | Left Middle Occipital Gyrus |
| | | | | Negative clusters | | |

| | | | | | | |
|---|---|---|---|---|---|---|
| 1 | 58.5 | -46.5 | 43.5 | -10.919 | 14229 | Right Inferior Parietal Lobule |
| 1a | 46.5 | -58.5 | 49.5 | -5.712 | | Right Angular Gyrus |
| 2 | -43.5 | -7.5 | -1.5 | -9.709 | 10179 | Left Insula Lobe |
| 2a | -61.5 | -19.5 | 7.5 | -5.636 | | Left Superior Temporal Gyrus |
| 3 | 4.5 | -28.5 | 46.5 | -8.396 | 5751 | Right Middle Cingulate Cortex |
| 3a | 13.5 | -43.5 | 46.5 | -4.368 | | Right Precuneus |
| 3b | 1.5 | -28.5 | 22.5 | -4.327 | | Left Posterior Cingulate Cortex |
| 4 | 43.5 | 49.5 | -1.5 | -7.495 | 5022 | Right Middle Orbital Gyrus |
| 4a | 34.5 | 58.5 | 16.5 | -5.291 | | Right Middle Frontal Gyrus |
| 4b | 46.5 | 46.5 | 19.5 | -5.146 | | Right Middle Frontal Gyrus |
| 4c | 34.5 | 55.5 | 28.5 | -4.754 | | Right Middle Frontal Gyrus |
| 5 | 43.5 | -4.5 | -1.5 | -6.756 | 3186 | Right Insula Lobe |
| 5a | 40.5 | -7.5 | -13.5 | -4.850 | | Right Hippocampus |
| 6 | 58.5 | -67.5 | -7.5 | -6.082 | 1242 | Right Inferior Temporal Gyrus |
| 6a | 49.5 | -52.5 | -1.5 | -4.733 | | Right Middle Temporal Gyrus |
| 7 | 13.5 | 34.5 | -13.5 | -5.988 | 1269 | Right Mid Orbital Gyrus |
| 7a | 10.5 | 25.5 | -4.5 | -4.158 | | Right Anterior Cingulate Cortex |
| 8 | -67.5 | -28.5 | 31.5 | -5.828 | 3618 | Left SupraMarginal Gyrus |
| 8a | -64.5 | -40.5 | 43.5 | -5.193 | | Left Inferior Parietal Lobule |
| 8b | -55.5 | -52.5 | 43.5 | -4.657 | | Left Inferior Parietal Lobule |
| 8c | -58.5 | -34.5 | 25.5 | -4.400 | | Left SupraMarginal Gyrus |
| 9 | 10.5 | -64.5 | 67.5 | -5.268 | 2943 | Right Precuneus |
| 9a | 13.5 | -79.5 | 49.5 | -5.050 | | Right Precuneus |
| 9b | 16.5 | -58.5 | 43.5 | -4.921 | | Right Precuneus |
| 9c | 16.5 | -76.5 | 61.5 | -4.721 | | Right Superior Parietal Lobule |
| 10 | -34.5 | -70.5 | 10.5 | -5.175 | 729 | Left Middle Occipital Gyrus |
| 11 | 37.5 | -61.5 | 4.5 | -5.120 | 999 | Right Middle Temporal Gyrus |
| 12 | 4.5 | 37.5 | 34.5 | -4.588 | 1053 | Right Middle Cingulate Cortex |
| 12a | 7.5 | 31.5 | 49.5 | -4.532 | | Right Superior Medial Gyrus |
| 13 | 31.5 | 34.5 | 43.5 | -4.563 | 864 | Right Middle Frontal Gyrus |

**Supplementary Table S6.** Peak coordinates of significant clusters of brain activation contrasting rumination and worry states. Peak points at least 12 mm apart were extracted from each cluster.

| Cluster ID | X | Y | Z | Peak Stat | Cluster Size ($mm^3$) | Area |
|---|---|---|---|---|---|---|
| | | | | Positive clusters | | |
| 1 | -64.5 | -25.5 | 43.5 | 5.097 | 837 | Left SupraMarginal Gyrus |
| 1a | -49.5 | -25.5 | 40.5 | 4.531 | | Left Inferior Parietal Lobule |
| 2 | -37.5 | -40.5 | 43.5 | 4.533 | 1269 | Left Inferior Parietal Lobule |

| | | | | | | |
|---|---|---|---|---|---|---|
| 2a | -46.5 | -37.5 | 55.5 | 3.945 | | Left Inferior Parietal Lobule |
| | | | Negative clusters | | | |
| 1 | 13.5 | -73.5 | 4.5 | -7.009 | 18279 | Right Calcarine Gyrus |
| 1a | -4.5 | -91.5 | 1.5 | -5.575 | | Left Calcarine Gyrus |
| 1b | 10.5 | -67.5 | -7.5 | -5.307 | | Right Lingual Gyrus |
| 1c | -31.5 | -76.5 | -13.5 | -5.069 | | Left Fusiform Gyrus |
| 2 | 25.5 | 31.5 | 58.5 | -6.815 | 27108 | Right Superior Frontal Gyrus |
| 2a | 34.5 | 19.5 | 40.5 | -6.702 | | Right Middle Frontal Gyrus |
| 2b | -1.5 | 43.5 | -10.5 | -6.625 | | Left Mid Orbital Gyrus |
| 2c | 19.5 | 55.5 | 25.5 | -6.110 | | Right Superior Frontal Gyrus |
| 3 | -22.5 | 25.5 | 61.5 | -6.814 | 8991 | Left Middle Frontal Gyrus |
| 3a | -10.5 | 13.5 | 67.5 | -6.695 | | Left SMA |
| 3b | -22.5 | 46.5 | 43.5 | -5.565 | | Left Superior Frontal Gyrus |
| 3c | -10.5 | 55.5 | 43.5 | -4.979 | | Left Superior Frontal Gyrus |
| 4 | 58.5 | -58.5 | 40.5 | -6.436 | 5697 | Right Inferior Parietal Lobule |
| 4a | 49.5 | -61.5 | 52.5 | -6.342 | | Right Angular Gyrus |
| 5 | -52.5 | -67.5 | 43.5 | -5.049 | 945 | Left Angular Gyrus |
| 6 | -16.5 | 52.5 | -1.5 | -4.996 | 810 | Left Superior Medial Gyrus |
| 7 | -7.5 | -91.5 | 31.5 | -4.898 | 864 | Left Cuneus |
| 7a | -19.5 | -88.5 | 31.5 | -4.714 | | Left Superior Occipital Gyrus |
| 8 | 1.5 | -28.5 | 34.5 | -4.830 | 1053 | Left Middle Cingulate Cortex |
| 8a | -4.5 | -19.5 | 40.5 | -3.838 | | Left Middle Cingulate Cortex |

**Supplementary Table S7.** Peak coordinates of significant clusters of brain activation contrasting rumination and positive thinking states. Peak points at least 12 mm apart were extracted from each cluster.

| Cluster ID | X | Y | Z | Peak Stat | Cluster Size ($mm^3$) | Area |
|---|---|---|---|---|---|---|
| | | | | | Positive clusters | |
| 1 | 28.5 | 22.5 | -13.5 | 6.193 | 918 | Right Insula Lobe |
| 1a | 40.5 | 25.5 | -4.5 | 3.640 | | Right Insula Lobe |
| 2 | 52.5 | -19.5 | -4.5 | 6.190 | 6669 | Right Superior Temporal Gyrus |
| 2a | 61.5 | -28.5 | -1.5 | 5.899 | | Right Middle Temporal Gyrus |
| 2b | 46.5 | -22.5 | -10.5 | 5.398 | | Right Middle Temporal Gyrus |
| 2c | 52.5 | -37.5 | -1.5 | 5.380 | | Right Middle Temporal Gyrus |
| 3 | 46.5 | 1.5 | -34.5 | 5.712 | 756 | Right Inferior Temporal Gyrus |
| 4 | -55.5 | -22.5 | -1.5 | 5.661 | 2619 | Left Middle Temporal Gyrus |
| 5 | -37.5 | 25.5 | -7.5 | 5.404 | 1350 | Left Inferior Frontal Gyrus (p. Orbitalis) |
| 5a | -49.5 | 22.5 | -4.5 | 4.415 | | Left Inferior Frontal Gyrus (p. Orbitalis) |
| 6 | -4.5 | 25.5 | 61.5 | 5.349 | 675 | Left Superior Medial Gyrus |

| | | | | | | |
|---|---|---|---|---|---|---|
| 7 | -49.5 | -1.5 | 1.5 | 5.323 | 972 | Left Superior Temporal Gyrus |
| 7a | -58.5 | 10.5 | 4.5 | 4.712 | | Left Rolandic Operculum |
| 8 | -58.5 | -64.5 | 31.5 | 5.077 | 1512 | Left Angular Gyrus |
| 8a | -58.5 | -52.5 | 28.5 | 4.433 | | Left SupraMarginal Gyrus |
| 8b | -40.5 | -52.5 | 28.5 | 4.082 | | Left Angular Gyrus |
| 9 | 10.5 | 16.5 | 61.5 | 5.037 | 810 | Right SMA |
| 9a | 13.5 | 28.5 | 58.5 | 4.517 | | Right Superior Medial Gyrus |
| 10 | -34.5 | 58.5 | 19.5 | 5.015 | 729 | Left Middle Frontal Gyrus |
| 10a | -22.5 | 64.5 | 19.5 | 4.586 | | Left Superior Frontal Gyrus |
| 10b | -34.5 | 61.5 | 13.5 | 4.431 | | Left Middle Frontal Gyrus |
| 11 | 49.5 | -25.5 | 43.5 | 4.779 | 729 | Right Postcentral Gyrus |
| 12 | -55.5 | -22.5 | 46.5 | 4.507 | 837 | Left Inferior Parietal Lobule |
| | | | | Negative clusters | | |
| 1 | 19.5 | 25.5 | 43.5 | -7.681 | 3402 | Right Superior Frontal Gyrus |
| 2 | 10.5 | -49.5 | 13.5 | -7.623 | 11367 | Right Precuneus |
| 2a | 7.5 | -58.5 | 22.5 | -6.464 | | Right Precuneus |
| 2b | -19.5 | -58.5 | 25.5 | -6.302 | | Left Cuneus |
| 2c | -10.5 | -52.5 | 13.5 | -6.112 | | Left Precuneus |
| 3 | 40.5 | -67.5 | 28.5 | -7.398 | 2916 | Right Middle Occipital Gyrus |
| 3a | 40.5 | -79.5 | 37.5 | -5.572 | | Right Middle Occipital Gyrus |
| 4 | 10.5 | 55.5 | -7.5 | -7.385 | 10692 | Right Mid Orbital Gyrus |
| 4a | -13.5 | 46.5 | -7.5 | -6.779 | | Left Mid Orbital Gyrus |
| 4b | -4.5 | 37.5 | 7.5 | -5.613 | | Left Anterior Cingulate Cortex |
| 4c | 7.5 | 37.5 | 4.5 | -4.811 | | Right Anterior Cingulate Cortex |
| 5 | -28.5 | -31.5 | -16.5 | -7.007 | 7614 | Left ParaHippocampal Gyrus |
| 5a | -22.5 | -13.5 | -16.5 | -5.744 | | Left Hippocampus |
| 5b | -31.5 | -40.5 | -4.5 | -5.351 | | Left ParaHippocampal Gyrus |
| 5c | -37.5 | -19.5 | -16.5 | -4.750 | | Left Fusiform Gyrus |
| 6 | -46.5 | 19.5 | -31.5 | -6.959 | 4050 | Left Medial Temporal Pole |
| 6a | -34.5 | 22.5 | -34.5 | -6.220 | | Left Medial Temporal Pole |
| 6b | -49.5 | 13.5 | -19.5 | -5.533 | | Left Temporal Pole |
| 6c | -49.5 | 13.5 | -40.5 | -5.485 | | Left Medial Temporal Pole |
| 7 | -40.5 | -76.5 | 34.5 | -6.503 | 2592 | Left Middle Occipital Gyrus |
| 7a | -31.5 | -82.5 | 46.5 | -4.729 | | Left Inferior Parietal Lobule |
| 8 | 16.5 | -79.5 | 1.5 | -6.497 | 3807 | Right Lingual Gyrus |
| 8a | 10.5 | -82.5 | 16.5 | -4.278 | | Right Cuneus |
| 9 | -25.5 | 1.5 | -31.5 | -6.489 | 1188 | Left ParaHippocampal Gyrus |
| 9a | -28.5 | 1.5 | -46.5 | -4.658 | | Left Fusiform Gyrus |
| 10 | -19.5 | 31.5 | 40.5 | -6.176 | 4752 | Left Middle Frontal Gyrus |
| 10a | -25.5 | 28.5 | 58.5 | -4.659 | | Left Middle Frontal Gyrus |

| | | | | | | |
|---|---|---|---|---|---|---|
| 11 | 40.5 | 22.5 | -34.5 | -6.082 | 2727 | Right Medial Temporal Pole |
| 11a | 43.5 | 13.5 | -43.5 | -5.510 | | Right Medial Temporal Pole |
| 11b | 31.5 | 10.5 | -34.5 | -5.104 | | Right Medial Temporal Pole |
| 11c | 31.5 | 25.5 | -37.5 | -4.527 | | Right Medial Temporal Pole |
| 12 | 28.5 | 37.5 | -7.5 | -5.975 | 1080 | Right Inferior Frontal Gyrus (p. Orbitalis) |
| 13 | -55.5 | -46.5 | -7.5 | -5.801 | 1701 | Left Middle Temporal Gyrus |
| 14 | 22.5 | -31.5 | -16.5 | -5.737 | 2997 | Right Fusiform Gyrus |
| 14a | 40.5 | -31.5 | -19.5 | -5.225 | | Right Fusiform Gyrus |
| 14b | 16.5 | -13.5 | -19.5 | -5.075 | | Right ParaHippocampal Gyrus |
| 14c | 34.5 | -37.5 | -10.5 | -4.728 | | Right ParaHippocampal Gyrus |
| 15 | -64.5 | -4.5 | -16.5 | -5.681 | 756 | Left Middle Temporal Gyrus |
| 15a | -67.5 | -16.5 | -10.5 | -5.340 | | Left Middle Temporal Gyrus |
| 16 | -34.5 | 34.5 | -10.5 | -5.591 | 972 | Left Inferior Frontal Gyrus (p. Orbitalis) |
| 17 | 25.5 | -4.5 | -46.5 | -5.587 | 1215 | Right Fusiform Gyrus |
| 17a | 28.5 | 4.5 | -34.5 | -3.932 | | Right Medial Temporal Pole |
| 18 | -10.5 | -88.5 | 1.5 | -5.172 | 3186 | Left Calcarine Gyrus |
| 19 | 43.5 | 31.5 | 13.5 | -5.164 | 810 | Right Inferior Frontal Gyrus (p. Triangularis) |

**Supplementary Table S8.** Peak coordinates of significant clusters of brain activation contrasting worry and positive thinking states. Peak points at least 12 mm apart were extracted from each cluster.

| Cluster ID | X | Y | Z | Peak Stat | Cluster Size ($mm^3$) | Area |
|---|---|---|---|---|---|---|
| | | | | | Positive clusters | |
| 1 | 13.5 | 25.5 | 55.5 | 8.430 | 21843 | Right SMA |
| 1a | -10.5 | 16.5 | 67.5 | 8.352 | | Left SMA |
| 1b | 7.5 | 37.5 | 55.5 | 6.659 | | Right Superior Medial Gyrus |
| 1c | -7.5 | 34.5 | 61.5 | 6.644 | | Left Superior Medial Gyrus |
| 2 | -46.5 | -28.5 | -4.5 | 7.401 | 8559 | Left Middle Temporal Gyrus |
| 2a | -55.5 | -37.5 | 7.5 | 5.952 | | Left Middle Temporal Gyrus |
| 2b | -67.5 | -43.5 | 4.5 | 5.025 | | Left Middle Temporal Gyrus |
| 2c | -46.5 | -46.5 | 1.5 | 4.916 | | Left Middle Temporal Gyrus |
| 3 | 55.5 | -52.5 | 43.5 | 7.359 | 6507 | Right Inferior Parietal Lobule |
| 3a | 46.5 | -58.5 | 49.5 | 6.088 | | Right Angular Gyrus |
| 4 | -43.5 | -1.5 | -37.5 | 7.279 | 1296 | Left Inferior Temporal Gyrus |
| 5 | -58.5 | 25.5 | 10.5 | 7.147 | 11529 | Left Inferior Frontal Gyrus (p. Triangularis) |
| 5a | -58.5 | 19.5 | 22.5 | 6.008 | | Left Inferior Frontal Gyrus (p. Triangularis) |
| 5b | -43.5 | 22.5 | -7.5 | 5.940 | | Left Inferior Frontal Gyrus (p. Orbitalis) |
| 5c | -37.5 | 55.5 | -7.5 | 5.578 | | Left Middle Orbital Gyrus |
| 6 | 46.5 | -1.5 | -34.5 | 6.838 | 1998 | Right Inferior Temporal Gyrus |

| | | | | | | |
|---|---|---|---|---|---|---|
| 7 | 49.5 | 40.5 | -7.5 | 6.571 | 6210 | Right Inferior Frontal Gyrus (p. Orbitalis) |
| 7a | 40.5 | 52.5 | -7.5 | 5.604 | | Right Middle Orbital Gyrus |
| 7b | 52.5 | 22.5 | 10.5 | 5.088 | | Right Inferior Frontal Gyrus (p. Triangularis) |
| 7c | 37.5 | 22.5 | -10.5 | 4.963 | | Right Inferior Frontal Gyrus (p. Orbitalis) |
| 8 | 64.5 | -34.5 | 1.5 | 6.454 | 6723 | Right Middle Temporal Gyrus |
| 8a | 43.5 | -40.5 | 1.5 | 4.864 | | Right Middle Temporal Gyrus |
| 9 | -64.5 | -52.5 | 28.5 | 6.379 | 6885 | Left SupraMarginal Gyrus |
| 9a | -52.5 | -61.5 | 37.5 | 6.362 | | Left Angular Gyrus |
| 9b | -43.5 | -52.5 | 31.5 | 5.265 | | Left Angular Gyrus |
| 10 | -1.5 | -16.5 | 40.5 | 6.262 | 3051 | Left Middle Cingulate Cortex |
| 11 | 40.5 | 25.5 | 49.5 | 6.227 | 2862 | Right Middle Frontal Gyrus |
| 11a | 49.5 | 16.5 | 49.5 | 4.443 | | Right Middle Frontal Gyrus |
| 12 | -16.5 | 46.5 | 40.5 | 5.916 | 7533 | Left Superior Frontal Gyrus |
| 12a | -25.5 | 52.5 | 25.5 | 5.639 | | Left Middle Frontal Gyrus |
| 12b | -37.5 | 58.5 | 13.5 | 5.273 | | Left Middle Frontal Gyrus |
| 12c | -28.5 | 67.5 | 7.5 | 5.237 | | Left Superior Frontal Gyrus |
| 13 | -46.5 | 19.5 | 46.5 | 5.863 | 1944 | Left Middle Frontal Gyrus |
| 14 | -25.5 | -73.5 | -28.5 | 4.985 | 675 | Left Cerebellum (Crus 1) |
| | | | | Negative clusters | | |
| 1 | 16.5 | -52.5 | 22.5 | -9.425 | 11259 | Right Precuneus |
| 1a | -16.5 | -55.5 | 19.5 | -8.793 | | Left Cuneus |
| 1b | 7.5 | -52.5 | 10.5 | -8.004 | | Right Calcarine Gyrus |
| 1c | -19.5 | -58.5 | 25.5 | -7.826 | | Left Cuneus |
| 2 | -34.5 | -34.5 | -13.5 | -8.528 | 4374 | Left Inferior Temporal Gyrus |
| 3 | -37.5 | -79.5 | 31.5 | -7.385 | 3591 | Left Middle Occipital Gyrus |
| 3a | -31.5 | -82.5 | 46.5 | -5.103 | | Left Inferior Parietal Lobule |
| 4 | -31.5 | 34.5 | -10.5 | -7.179 | 1296 | Left Inferior Frontal Gyrus (p. Orbitalis) |
| 5 | -31.5 | -1.5 | -31.5 | -6.598 | 7344 | Left Fusiform Gyrus |
| 5a | -22.5 | -13.5 | -16.5 | -6.275 | | Left Hippocampus |
| 5b | -46.5 | 19.5 | -28.5 | -5.946 | | Left Medial Temporal Pole |
| 5c | -31.5 | -1.5 | -13.5 | -5.399 | | Left Amygdala |
| 6 | -58.5 | 1.5 | -19.5 | -6.312 | 1242 | Left Middle Temporal Gyrus |
| 6a | -67.5 | -16.5 | -10.5 | -4.797 | | Left Middle Temporal Gyrus |
| 7 | 43.5 | -70.5 | 28.5 | -5.894 | 2511 | Right Middle Occipital Gyrus |
| 7a | 37.5 | -79.5 | 40.5 | -5.118 | | Right Middle Occipital Gyrus |
| 7b | 46.5 | -79.5 | 28.5 | -4.523 | | Right Middle Occipital Gyrus |
| 8 | 19.5 | -13.5 | -16.5 | -5.873 | 1350 | Right Hippocampus |
| 8a | 37.5 | -13.5 | -16.5 | -4.123 | | Right Hippocampus |
| 9 | 43.5 | 19.5 | -31.5 | -5.804 | 783 | Right Temporal Pole |
| 10 | 28.5 | 37.5 | -7.5 | -5.763 | 675 | Right Inferior Frontal Gyrus (p. Orbitalis) |

| | | | | | | |
|---|---|---|---|---|---|---|
| 11 | 16.5 | -34.5 | -10.5 | -5.052 | 1242 | Right Lingual Gyrus |
| 11a | 28.5 | -31.5 | -16.5 | -4.761 | | Right Fusiform Gyrus |
| 11b | 37.5 | -22.5 | -13.5 | -4.036 | | Right Hippocampus |

**Supplementary Table S9.** Peak coordinates of significant clusters of brain activation showing age effect during the rumination state. Peak points at least 12 mm apart were extracted from each cluster.

| Cluster ID | X | Y | Z | Peak Stat | Cluster Size ($mm^3$) | Area |
|---|---|---|---|---|---|---|
| | | | | Positive clusters | | |
| 1 | 46.5 | 46.5 | 13.5 | 5.477 | 2403 | Right Middle Frontal Gyrus |
| 1a | 37.5 | 58.5 | 19.5 | 3.916 | | Right Middle Frontal Gyrus |
| 2 | 4.5 | -79.5 | -7.5 | 5.309 | 7668 | Right Lingual Gyrus |
| 2a | -4.5 | -76.5 | -25.5 | 5.162 | | Left Cerebellum (Crus 1) |
| 2b | -19.5 | -82.5 | -25.5 | 4.817 | | Left Cerebellum (Crus 1) |
| 2c | 4.5 | -91.5 | -10.5 | 4.577 | | Left Calcarine Gyrus |
| 3 | 13.5 | -52.5 | -25.5 | 5.070 | 4887 | Right Cerebellum (IV V) |
| 3a | 34.5 | -49.5 | -19.5 | 4.815 | | Right Fusiform Gyrus |
| 3b | 16.5 | -52.5 | -13.5 | 4.724 | | Right Cerebellum (IV V) |
| 3c | 25.5 | -55.5 | -31.5 | 4.249 | | Right Cerebellum (VI) |
| 4 | 25.5 | 13.5 | 67.5 | 4.827 | 1998 | Right Superior Frontal Gyrus |
| 4a | 40.5 | 10.5 | 61.5 | 4.579 | | Right Middle Frontal Gyrus |
| 4b | 28.5 | 7.5 | 64.5 | 4.331 | | Right Superior Frontal Gyrus |
| 4c | 31.5 | 7.5 | 52.5 | 3.976 | | Right Middle Frontal Gyrus |
| 5 | 49.5 | -37.5 | 34.5 | 4.822 | 2619 | Right SupraMarginal Gyrus |
| 5a | 46.5 | -43.5 | 61.5 | 4.345 | | Right Superior Parietal Lobule |
| 5b | 46.5 | -40.5 | 46.5 | 3.567 | | Right Inferior Parietal Lobule |
| 6 | 16.5 | -10.5 | 19.5 | 4.810 | 1701 | Right Caudate Nucleus |
| 7 | -37.5 | -73.5 | -16.5 | 4.491 | 1350 | Left Fusiform Gyrus |
| 7a | -31.5 | -70.5 | -28.5 | 4.277 | | Left Cerebellum (Crus 1) |
| 7b | -37.5 | -79.5 | -16.5 | 4.144 | | Left Fusiform Gyrus |
| 8 | 7.5 | 31.5 | 43.5 | 4.424 | 1377 | Right Superior Medial Gyrus |
| 9 | -34.5 | -16.5 | 73.5 | 4.392 | 1620 | Left Precentral Gyrus |
| 9a | -43.5 | -19.5 | 64.5 | 4.144 | | Left Precentral Gyrus |
| 10 | 7.5 | -76.5 | 58.5 | 4.376 | 1890 | Right Precuneus |
| 10a | -4.5 | -70.5 | 61.5 | 4.228 | | Left Precuneus |
| 10b | -19.5 | -73.5 | 61.5 | 4.198 | | Left Superior Parietal Lobule |
| 10c | -10.5 | -67.5 | 55.5 | 4.102 | | Left Precuneus |

**Supplementary Table S10.** Peak coordinates of significant clusters of brain activation showing age effect during the worry state. Peak points at least 12 mm apart were extracted from each cluster.

| Cluster ID | X | Y | Z | Peak Stat | Cluster Size (mm$^3$) | Area |
|---|---|---|---|---|---|---|
| | | | | Negative clusters | | |
| 1 | 55.5 | -61.5 | 13.5 | -5.063 | 1809 | Right Middle Temporal Gyrus |
| 1a | 52.5 | -46.5 | 13.5 | -4.292 | | Right Superior Temporal Gyrus |
| 2 | -1.5 | -7.5 | 37.5 | -4.604 | 1674 | Left Middle Cingulate Cortex |
| 2a | -1.5 | -16.5 | 46.5 | -4.007 | | Left Middle Cingulate Cortex |
| 2b | -10.5 | -4.5 | 37.5 | -3.569 | | Left Middle Cingulate Cortex |
| 3 | 1.5 | 22.5 | 28.5 | -4.31 | 1080 | Left Anterior Cingulate Cortex |
| 3a | 4.5 | 31.5 | 19.5 | -3.739 | | Right Anterior Cingulate Cortex |

**Supplementary Table S11.** Peak coordinates of significant clusters of brain activation contrasting age effects between rumination and worry states. Peak points at least 12 mm apart were extracted from each cluster.

| Cluster ID | X | Y | Z | Peak Stat | Cluster Size (mm$^3$) | Area |
|---|---|---|---|---|---|---|
| | | | | Positive clusters | | |
| 1 | 55.5 | -61.5 | 13.5 | 6.357 | 17847 | Right Middle Temporal Gyrus |
| 1a | 49.5 | -52.5 | 1.5 | 5.766 | | Right Middle Temporal Gyrus |
| 1b | 46.5 | -40.5 | 55.5 | 5.715 | | Right Inferior Parietal Lobule |
| 1c | 55.5 | -40.5 | 31.5 | 5.562 | | Right SupraMarginal Gyrus |
| 2 | -37.5 | -67.5 | 13.5 | 6.043 | 16335 | Left Middle Temporal Gyrus |
| 2a | -40.5 | -85.5 | 22.5 | 5.590 | | Left Middle Occipital Gyrus |
| 2b | -31.5 | -88.5 | 31.5 | 5.560 | | Left Middle Occipital Gyrus |
| 2c | -52.5 | -58.5 | 1.5 | 5.355 | | Left Middle Temporal Gyrus |
| 3 | -34.5 | 28.5 | 31.5 | 5.680 | 5967 | Left Middle Frontal Gyrus |
| 3a | -31.5 | 46.5 | 16.5 | 5.441 | | Left Middle Frontal Gyrus |
| 3b | -34.5 | 34.5 | 19.5 | 5.030 | | Left Middle Frontal Gyrus |
| 3c | -40.5 | 43.5 | 34.5 | 4.888 | | Left Middle Frontal Gyrus |
| 4 | 40.5 | -79.5 | 31.5 | 5.473 | 2943 | Right Middle Occipital Gyrus |
| 4a | 46.5 | -73.5 | 43.5 | 5.207 | | Right Angular Gyrus |
| 4b | 37.5 | -70.5 | 25.5 | 4.748 | | Right Middle Occipital Gyrus |
| 5 | -16.5 | -82.5 | -22.5 | 5.438 | 2862 | Left Cerebellum (Crus 1) |
| 5a | -4.5 | -64.5 | -28.5 | 4.115 | | Left Cerebellum (VIII) |
| 6 | -4.5 | -61.5 | 67.5 | 5.433 | 10449 | Left Precuneus |
| 6a | 1.5 | -19.5 | 46.5 | 5.389 | | Left Middle Cingulate Cortex |
| 6b | -10.5 | -49.5 | 55.5 | 5.051 | | Left Precuneus |
| 6c | 10.5 | -40.5 | 61.5 | 4.835 | | Right Paracentral Lobule |

| | | | | | | |
|---|---|---|---|---|---|---|
| 7 | 10.5 | 28.5 | 22.5 | 5.354 | 1134 | Right Anterior Cingulate Cortex |
| 8 | 28.5 | 10.5 | 64.5 | 5.313 | 3078 | Right Superior Frontal Gyrus |
| 8a | 25.5 | 13.5 | 52.5 | 4.324 | | Right Middle Frontal Gyrus |
| 9 | 31.5 | 43.5 | 16.5 | 5.229 | 4860 | Right Middle Frontal Gyrus |
| 9a | 43.5 | 40.5 | 13.5 | 4.897 | | Right Middle Frontal Gyrus |
| 9b | 31.5 | 37.5 | 19.5 | 4.698 | | Right Middle Frontal Gyrus |
| 9c | 40.5 | 31.5 | 13.5 | 4.425 | | Right Inferior Frontal Gyrus (p. Triangularis) |
| 10 | 49.5 | 16.5 | 31.5 | 5.183 | 4482 | Right Inferior Frontal Gyrus (p. Opercularis) |
| 10a | 52.5 | 1.5 | 37.5 | 5.086 | | Right Precentral Gyrus |
| 10b | 40.5 | 10.5 | 40.5 | 4.412 | | Right Middle Frontal Gyrus |
| 10c | 37.5 | -1.5 | 37.5 | 4.374 | | Right Precentral Gyrus |
| 11 | 10.5 | -31.5 | -25.5 | 5.167 | 7992 | Right Cerebellum (III) |
| 11a | 16.5 | -46.5 | -13.5 | 5.051 | | Right Cerebellum (IV V) |
| 11b | 37.5 | -52.5 | -31.5 | 5.033 | | Right Cerebellum (Crus 1) |
| 11c | 22.5 | -43.5 | -28.5 | 4.943 | | Right Cerebellum (IV V) |
| 12 | 19.5 | -55.5 | 55.5 | 5.148 | 2700 | Right Superior Parietal Lobule |
| 12a | 13.5 | -70.5 | 52.5 | 4.696 | | Right Superior Parietal Lobule |
| 12b | 25.5 | -70.5 | 61.5 | 4.509 | | Right Superior Parietal Lobule |
| 13 | -43.5 | 55.5 | -7.5 | 5.145 | 891 | Left Middle Orbital Gyrus |
| 13a | -25.5 | 49.5 | -4.5 | 4.119 | | Left Middle Orbital Gyrus |
| 14 | 7.5 | -1.5 | 37.5 | 5.031 | 972 | Right Middle Cingulate Cortex |
| 15 | -28.5 | 4.5 | 58.5 | 4.727 | 891 | Left Middle Frontal Gyrus |
| 16 | 40.5 | 25.5 | -4.5 | 4.678 | 1242 | Right Insula Lobe |
| 17 | 16.5 | -82.5 | -19.5 | 4.654 | 1377 | Right Cerebellum (Crus 1) |
| 17a | 28.5 | -79.5 | -19.5 | 4.594 | | Right Cerebellum (VI) |
| 17b | 10.5 | -73.5 | -13.5 | 4.182 | | Right Cerebellum (VI) |
| 18 | -28.5 | 22.5 | -7.5 | 4.601 | 945 | Left Insula Lobe |
| 19 | -37.5 | 13.5 | 22.5 | 4.419 | 1350 | Left Inferior Frontal Gyrus (p. Opercularis) |
| 19a | -46.5 | 7.5 | 28.5 | 4.022 | | Left Inferior Frontal Gyrus (p. Opercularis) |
| 20 | 1.5 | 37.5 | 49.5 | 4.416 | 2943 | Left Superior Medial Gyrus |
| 20a | 1.5 | 25.5 | 52.5 | 4.249 | | Left SMA |
| 20b | 1.5 | 49.5 | 37.5 | 4.166 | | Left Superior Medial Gyrus |
| 20c | 10.5 | 22.5 | 46.5 | 4.132 | | Right SMA |

**Supplementary Table S12.** Peak coordinates of significant clusters of brain activation contrasting age effects between rumination and positive thinking states. Peak points at least 12 mm apart were extracted from each cluster.

| Cluster ID | X | Y | Z | Peak Stat | Cluster Size ($mm^3$) | Area |
|---|---|---|---|---|---|---|
| | | | | Positive clusters | | |
| 1 | 55.5 | -43.5 | 40.5 | 5.111 | 1755 | Right SupraMarginal Gyrus |
| 2 | 49.5 | 16.5 | 22.5 | 4.933 | 2025 | Right Inferior Frontal Gyrus (p. Triangularis) |
| 2a | 43.5 | 4.5 | 28.5 | 3.771 | | Right Precentral Gyrus |

**Supplementary Table S13.** Peak coordinates of significant clusters of brain activation contrasting age effects between worry and positive thinking states. Peak points at least 12 mm apart were extracted from each cluster.

| Cluster ID | X | Y | Z | Peak Stat | Cluster Size ($mm^3$) | Area |
|---|---|---|---|---|---|---|
| | | | | Negative clusters | | |
| 1 | 40.5 | -82.5 | 28.5 | -5.334 | 1053 | Right Middle Occipital Gyrus |
| 2 | 52.5 | -7.5 | 4.5 | -5.059 | 891 | Right Heschls Gyrus |
| 3 | -19.5 | -40.5 | -7.5 | -4.941 | 891 | Left ParaHippocampal Gyrus |
| 3a | -22.5 | -52.5 | 7.5 | -4.044 | | Left Precuneus |
| 4 | -28.5 | 10.5 | 58.5 | -4.834 | 1593 | Left Middle Frontal Gyrus |

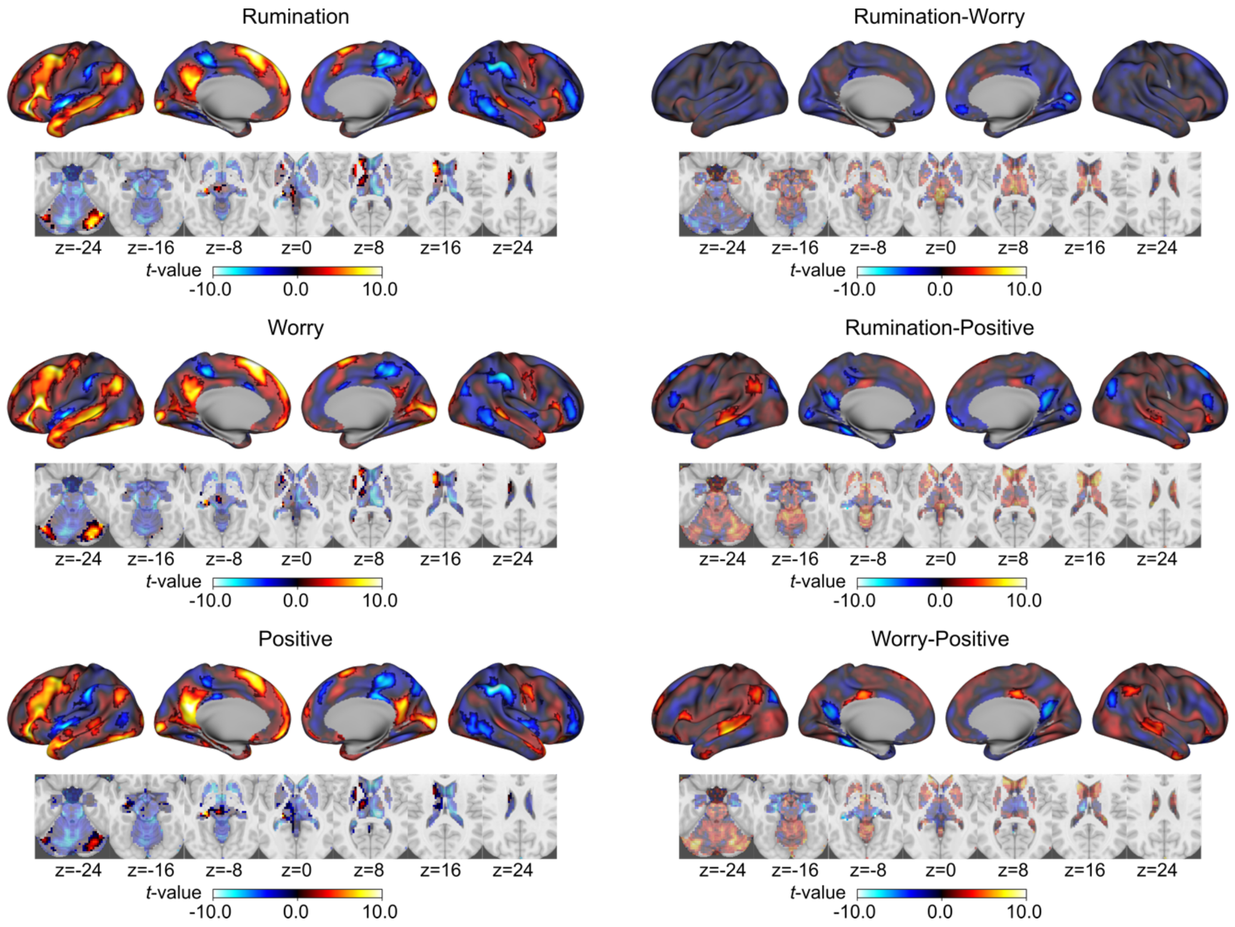

**Supplementary Figure S1**. Brain activation patterns associated with the thought states and their contrasts, evaluated using block-wise response models. The maps are displayed on the inflated cortical surface, while subcortical and cerebellar regions are shown on axial slice maps. Clusters with voxel-wise $p < 0.001$, corrected for cluster extent at $p < 0.05$, are highlighted with opaque colors.
